# Selection for Pollen Competitive Ability in Mixed Mating Systems

**DOI:** 10.1101/222117

## Abstract

Co-expression of genes in plant sporophytes and gametophytes allows correlated gametic and sporophytic selection. Theory predicts that, under outcrossing, an allele conferring greater pollen competitive ability should fix within a population unless antagonistic pleiotropy with the sporophyte stage is strong. However, under strong selfing, pollen competitiveness is immaterial as superior and inferior competitors are deposited on opposite stigmas, producing assortative competition. Because many plant species have mixed-mating systems, selfing should be critical in the spread and maintenance of pollen-expressed genes affecting competitiveness. We present two one-locus, two-allele population genetic models for the evolution of a locus controlling pleiotropic antagonism between pollen competitiveness and diploid fitness. Analytical solutions provide minimum and maximum selfing rates allowing invasion of alleles with greater diploid and haploid fitness respectively. Further, polymorphism is only maintained when diploid selection is recessive. Fixation of the allele conferring greater pollen competitiveness may be prevented, even with weak sporophytic counter-selection, with sufficiently high selfing. Finally, selfing expands and limits the range of haploid-diploid selection coefficients allowing polymorphism, depending on dominance and selfing mode.

## INTRODUCTION

Over 7000 genes are expressed in male gametophytes—pollen grains and tubes and the sperm they transmit—as they compete in the pistil for ovules (Page and Grossniklaus 2002; Honys and Twell 2003; Rutley and Twell 2015). Upwards of 60% of known plant genes are expressed in both the gametophyte and sporophyte (Borges et al. 2009; Arunkumar et al. 2013). For example, over 100 cases of co-expression in pollen tubes and root hairs have been identified in tobacco (Hadif et al. 2012). This high degree of co-expression presents opportunity for pollen selection to affect sporophyte evolution, and *vice versa*. Haldane (1932) recognized this potential, writing “Clearly a higher plant species is at the mercy of its pollen grains. A gene which greatly accelerates pollen tube growth will spread through a species even if it causes moderately disadvantageous changes in the adult plant.” Simply, genes conferring an advantage in the gametophyte should fix unless there is strong antagonistic pleiotropy with sporophyte fitness.

Selection should be strong on alleles with concordant impacts on both gametophyte and sporophyte stages. Deleterious recessive mutations, which are masked in heterozygous sporophytes, are fully exposed to selection in haploid gametophytes. This facilitates purging, diminishing inbreeding depression (Charlesworth and Charlesworth 1992). Some have suggested that alleles promoting pollen competition will increase offspring quality: large stigmatic pollen loads lead to intense competition in the style, favouring genetically superior gametophytes, which contribute their genes to the seed (Mulcahy and Mulcahy 1987; Winsor et al. 1987; Winsor et al. 2000; but see Baskin and Baskin 2015; Pélabon et al. 2016). Walsh and Charlesworth (1992) found that when fitness effects on the gametophyte and sporophyte are concordant, selection will quickly eliminate deleterious mutations and drive favourable ones to fixation. But, as Haldane suggested, polymorphism may be maintained if increased pollen competitive ability is counterbalanced by costs to the sporophyte.

Ploidally antagonistic selection has been considered previously (Ewing 1977). Immler, Arnqvist and Otto (2012) explored the maintenance of genetic variation under discordant selection between haploid and diploid phases, as well as under sex-specific selection.

Considering an autosomal gene, they demonstrated polymorphism is most easily maintained under negative ploidy-by-sex interactions, under which sex-specific selection acts in opposite directions in haploids and diploids. They conclude that polymorphism can be maintained, even in the absence of sex differences in selection, when selection is opposing between sporophytic and gametophytic stages.

High rates of self-pollination diminish the advantage of pollen competitiveness due to assortative competition (Mazer et al. 2010). Imagine a one locus system, with allele *C_g_* conferring a competitive advantage to pollen (i.e., in the gametophytic phase) and the alternate allele *C_s_* advantages in the sporophytic phase. When a *C_g_C_g_* homozygote self-pollinates, the *C_g_* allele gets passed to the offspring by necessity. On the opposite homozygote, the *C_s_* allele is likewise successful by necessity, despite its competitive inferiority. The competitive advantage of *C_g_*-over *C_s_*-bearing pollen is realized when heterozygotes self-pollinate, but because selfing reduces heterozygote frequency, over time there are increasingly fewer opportunities for *C_g_* to exercise its superiority. Thus, with obligate selfing, the *C_s_* allele gains refuge from competition and quickly goes to fixation due to its higher sporophytic fitness. Consistent with these predictions, Mazer et al. (2018) demonstrated faster pollen tube growth rates in the predominantly outcrossing *Clarkia unguiculate* (Onagraceae) compared to the facultatively selfing *C. exilis*. A similar result was found for selfing and outcrossing populations of *Clarkia tembloriensis* (Smith-Huerta 1996).

About 40% of plant species have mixed-mating systems: individuals receive mixtures of self and outcross pollen (Schemske and Lande 1985; Goodwillie et al. 2005). Of the 155 studies of mixed-mating systems in plants reviewed by Barrett and Eckert (2012), two-thirds report selfing rates in excess of 20%. In Whitehead et al. (2018), it was found that 63% of 105 surveyed species had at least one population with a mixed-mating system. How do intermediate selfing rates influence the fate of a new mutation increasing pollen competitiveness at the expense of the sporophyte?

An important paper by Jordan and Connallon (2014) considered the impact of self-fertilization on sexual conflict in hermaphrodites. They found that selfing narrows invasion conditions of new alleles conferring male-benefit in the haploid phase if they impose a cost on diploid female function. Their model considered the case in which gamete quality is determined by parental genotype. In this situation, a *C_g_*-bearing gamete from a homozygous adult differs from a *C_g_*-bearing gamete produced by a heterozygote, weakening the gamete haplotype-phenotype correspondence. Gene expression in animal sperm is limited (Parker and Begon 1993), and so it is reasonable to assume genetic paternal effects on sperm quality for these species. In plants, adult diploid condition can affect pollen size and quality (Young and Stanton 1990; Mazer and Gorchov 1996; Galloway 2001; Smith-Huerta et al. 2007). Pollen size, which could well be mediated by genetically influenced resource allocation in the sporophyte, readily responds to artificial selection (Sarkissian and Harder 2001; McCallum and Chang 2016).

We explore a different situation, in which pollen competitiveness is affected by genes expressed directly in the male gametophyte. Here one may expect stronger selection responses due to stronger haplotype-phenotype correspondence. In our one-locus model, allele *C_g_* increases pollen competitive ability at the expense of sporophyte fitness. Analytical solutions show there are maximum and minimum selfing rates allowing invasion of *C_g_* and *C_s_* alleles, respectively, and that stable polymorphism is possible only when *C_g_* is recessive. Further analyses show that as selfing rate increases, *C_g_*’s competitive advantage is negated by increasingly smaller diploid cost.

## METHODS

### The models

Departures from Hardy-Weinberg equilibrium via selfing necessitate that selection be modelled in terms of genotype frequencies (Crow and Kimura 1970). We refer to the frequencies of the *C_g_C_g_, C_g_C_s_* and *C_s_C_s_* genotypes as *X*_0_, *X*_1_ and *X*_2_, respectively, where subscripts denote the number of *C_s_* alleles. Allele frequencies are *p* and *q* for *C_g_* and *C_s_*, respectively.

Assume a pleiotropic effect by which the *C_g_* allele increases pollen competitive ability but decreases the number of pollen grains and ovules that the diploid plant contributes to the gamete pool, either by reduced survival or lower flower production. The *C_s_* allele is favoured through both sex functions in the diploid stage, disfavoured in haploid pollen and has no effect on ovules. Fitness of sporophyte *C_s_C_s_* is *W_2_* = 1, and that of *C_g_C_g_* is *W_0_* = 1−*s*, while the heterozygote *C_g_C_s_* is *W_1_* = 1−*hs*, where *h* is the degree of dominance for fitness. As selection may act either through survival or fertility, we assume a common dominance coefficient for male and female function. (For a discussion of the effects of sex-specific dominance, see Appendix A).

The absolute competitive ability of pollen bearing the *C_g_* allele is 1 and that of pollen bearing the *C_s_* allele is 1-*t*. All individual plants are expected to have many flowers. Population size is effectively infinite.

The level of self-pollination in the model depends on r, the proportion of pollen allocated to self-mating, which reflects the potential rate of self-fertilization. Absent selection, *r* equals the proportion of zygotes that are produced by self-pollination, *i.e*., the realized selfing rate. As shown below, potential and realized selfing rates can differ when selection acts on the gametophyte. For sake of brevity, *r* is hereafter referred to as selfing rate. We assume all plants have the same *r* value. We model two selfing forms: fixed selfing and mass-action selfing, defined below. The models we present assume haploid genetic control over pollen competitive ability. As we present our results, we will for comparison also present results using models by Jordan and Connallon (2014; see Appendix B), which assume diploid control.

### FIXED selfing

The fixed selfing model echoes the biology of species such as *Impatiens* and *Viola*, where cleistogamous flowers are obligate selfers, and chastogamous flowers outcross (Schemske 1978, Winn and Moriuchi 2009). In this model, proportion *r* of flowers on each plant is dedicated to selfing while 1−*r* of flowers is allocated to outcrossing. Thus, a stigma receives either only self-pollen or only outcross-pollen. Selection for pollen competitiveness occurs on all outcrossing flowers, but among selfing flowers, occurs only on heterozygotes. Our recursion equations for genotype frequencies consider selection during the diploid and haploid stages in separate steps. After diploid selection, diploid genotype frequencies are:

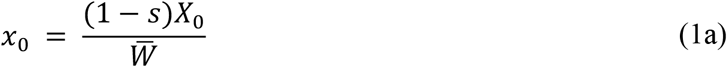

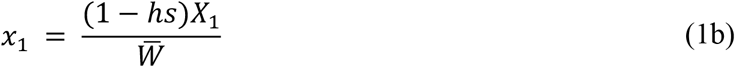

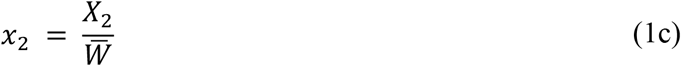

where 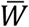 is the mean diploid fitness:

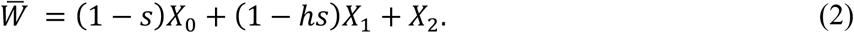

After mating and haploid selection, offspring genotype frequencies are

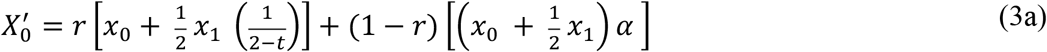

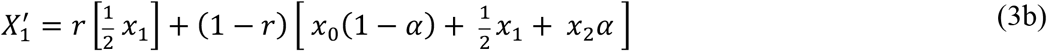

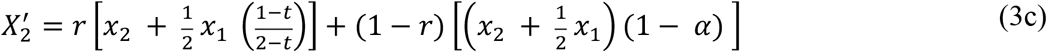

where *α* reflects the relative competitiveness of *C_g_*-bearing pollen as a function of the absolute competitiveness conferred by the *C_g_* allele and the frequency of the *C_g_* allele in the pollen pool as a result of genotype frequency following selection:

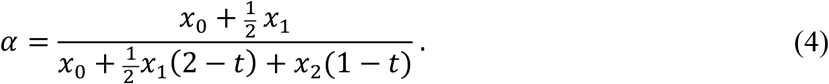

It follows that the relative competitiveness of *C_s_*-bearing pollen is (1−*α*).

To understand Equations (3a-c), consider separately the offspring produced on selfing (*r*) and outcrossing (1−*r*) flowers. Under selfing, competition among pollen tubes carrying opposite alleles only occurs within heterozygote pistils. Assuming heterozygotes produce *C_g_* and *C_s_* pollen in equal numbers, ½ of their offspring will likewise be heterozygous, with the remaining offspring divided between *C_g_C_g_* and *C_s_C_s_* homozygotes in proportions of 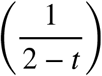 and 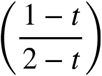 (Appendix C). We assume each selfing flower produces enough pollen to fertilise its ovules, such that only female diploid fitness terms, and not α (*i.e*., male diploid fitness terms), are included. Simply, the number of offspring produced via selfing is limited by seed production, not pollen production. Offspring produced through outcrossing will be a function of both female and male diploid fitness, as well as allelic differences in pollen competitive ability, specified by α. For simplicity, we make no provision for inbreeding depression; increasing failure rate of selfed offspring has the same effect as reducing the basic selfing rate (see Jordan and Connallon 2014).

### Mass-action selfing

Mass-action selfing (Holsinger 1991) echoes more general modes of mixed-mating. Here *r* still denotes the proportion of pollen individuals allocate to selfing, but in this case all stigmas receive a mix of self- and outcross-pollen, resulting in inter-allelic competition for all ovules. Under this scenario, the amount of self-pollen received by a stigma is a function of two factors, one being *r*, and the other being the quantity of pollen produced by the individual relative to that it receives through outcrossing. Thus, the true selfing rate is dependent on genotype frequency. We make no provisions for pollen discounting here but consider this elaboration in Appendix D.

Assuming diploid selection occurs according to Equations (1a-c), we arrive at the following genotype frequencies after mating and haploid selection:

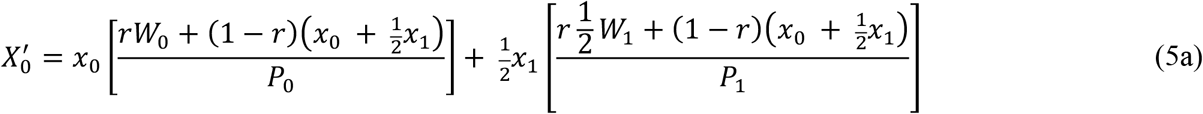

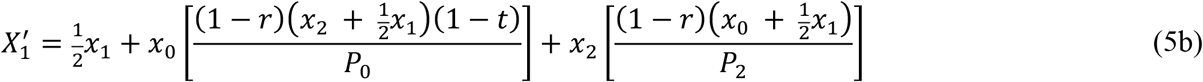

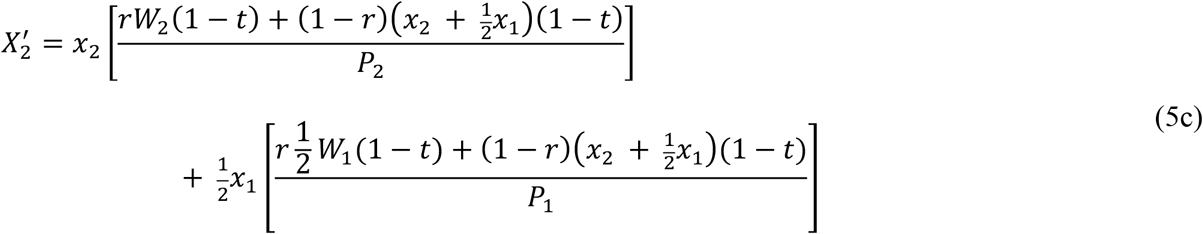

where *P_0_, P_1_* and *P_2_* give the total pollen received by *X_0_, X_1_* and *X*_2_ individuals respectively and are defined as:

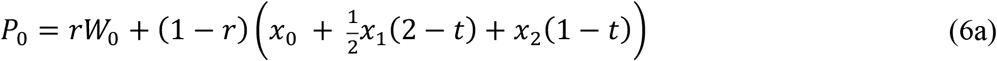

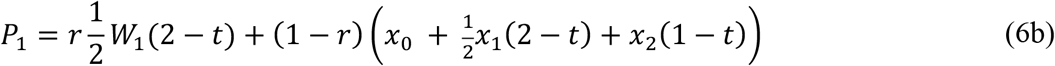

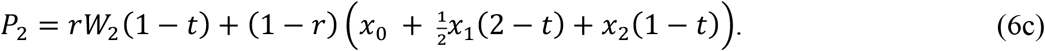

In Eq. 5a, the two terms represent *C_g_C_g_* offspring seed produced by *C_g_C_g_* and *C_g_C_s_* plants with post-diploid selection frequencies of *x*_0_ and *x*_1_. All ovules on the homozygote carry *C_g_*, whereas on heterozygotes only half do. The terms inside the square brackets are the probability that an ovule receives *C_g_*-bearing sperm, which depends on the amount of successful *C_g_* pollen (numerator) divided by the total amount of pollen arriving at the stigma (denominator). Eq. 5c follows similar logic for *C_s_C_s_* offspring. Turning to Eq. 5b, the three terms represent *C_g_C_s_* offspring produced by *C_g_C_s_*, *C_g_C_g_* and *C_s_C_s_* plants, respectively. Half of all seed produced by *C_g_C_s_* individuals are also heterozygotes. The proportion of heterozygotes produced by *C_g_C_g_* individuals are a function of the relative abundance and competitiveness of *C_s_* pollen (numerator) compared to the total pollen arriving on *C_g_C_g_* stigmas. This proportion is given by the term inside the first square bracket. *C_g_C_s_* offspring from *C_s_C_s_* individuals follow identical logic but for *C_g_* pollen.

### Software and computation

All analytical analyses were derived using Wolfram Mathematica 11.0.1.0. To find conditions for stable polymorphism, we derived leading eigenvalues for Jacobian Matrices using genotype recursions for gametophytic control (Eqs. 3a-c, 5a-c) and sporophytic control (Eqs. B9–B11, Appendix B) of pollen competitiveness and determined the conditions that allowed invasion at a boundary by solving for parameters producing a leading eigenvalue of absolute value greater than 1. Invasion analyses were performed for two boundaries: the *X*_0_ boundary, at which the *C_s_* allele appears in a population fixed for the *C_g_* allele (i.e.,*p* = 1), and the *X*_2_ boundary, at which the *C_g_* allele appears in a population fixed for the *C_s_* allele (i.e.,*p* = 0). Code for analytical analyses is provided in the online supplemental materials.

To find expressions for equilibrium allele frequencies, we generated power series expansions (assuming weak selection for simplification) for expressions giving the single-generation change in genotype frequencies. We then solved for the equilibrium by finding the *p* value yielding zero change. Predicted equilibrium frequencies were verified numerically. Numerical methods and results are presented in Appendix E.

## RESULTS

### Analytical invasion conditions

We explored the impact of selfing rate on invasion potential for both *C_g_* and *C_s_* when rare. For fixed selfing, analytical conditions for invasion at the *X*_0_ and *X*_2_ boundaries are derived using Equations (3a-c). Likewise, analytical conditions for mass-action selfing use Equations (5a-c). For comparison with diploid control models (Jordan and Connallon 2014) and, in the case of mass-action selfing, to produce tractable solutions, we assume weak selection (i.e., 0 < *s, t* ≪ 1), though we do find good agreement between invasion conditions predicted under fixed selfing both with and without weak selection (Appendix E).

### Fixed selfing

Under fixed selfing, boundary selfing rates permitting invasion of the *C_g_* and *C_s_* alleles, respectively, are as follows:

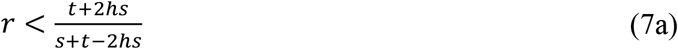

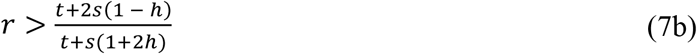

These analytical solutions are visualised in Figure (1a-c). Under random mating (i.e., *r* = 0), and when either fully dominant or recessive (*h* = 0 or 1), the *C_g_* allele invades and goes to fixation if *s/t* < 0.5 (*i.e*., when the haploid competitive benefit is more than twice the diploid fitness cost). When diploid fitness effects are additive (*h* = 0.5), fixation occurs even as *s/t* approaches 1.0. *C_g_* wins under random mating as the *C_s_* allele has no refuge from competition. Under complete selfing (*r* = 1), *C_g_* never invades. Pollen competition occurs only on heterozygotes, and since selfing reduces heterozygote frequencies, opportunities for *C_g_*-bearing pollen to outcompete *C_s_* effectively vanish. As *C_s_* confers higher sporophyte fitness in the homozygous state, it increases when rare.

The analytical solutions indicate that polymorphism can be maintained in mixed-mating systems (0 < *r* < 1) with fixed selfing only if the deleterious effect of the *C_g_* allele in the diploid phase is recessive (*h* < 0.5) (Figures 1a-c, 2). When completely recessive (*h* = 0), there is a range of selfing rates in which both alleles can invade, implying protected polymorphism. The *C_g_* allele invades because, when rare, it occurs exclusively in heterozygotes, where its negative effect on diploid fitness is masked from selection and moderate selfing allow opportunities to outcompete *C_s_*-bearing pollen for ovules on heterozygotes. Within this range of selfing rates, the *C_s_* allele also increases when rare because, 1) heterozygotes have higher relative fitness than *C_g_C_g_* homozygotes and 2) selfing reduces exposure of *C_s_* pollen to competition with superior *C_g_* pollen.

**Figure 2.**
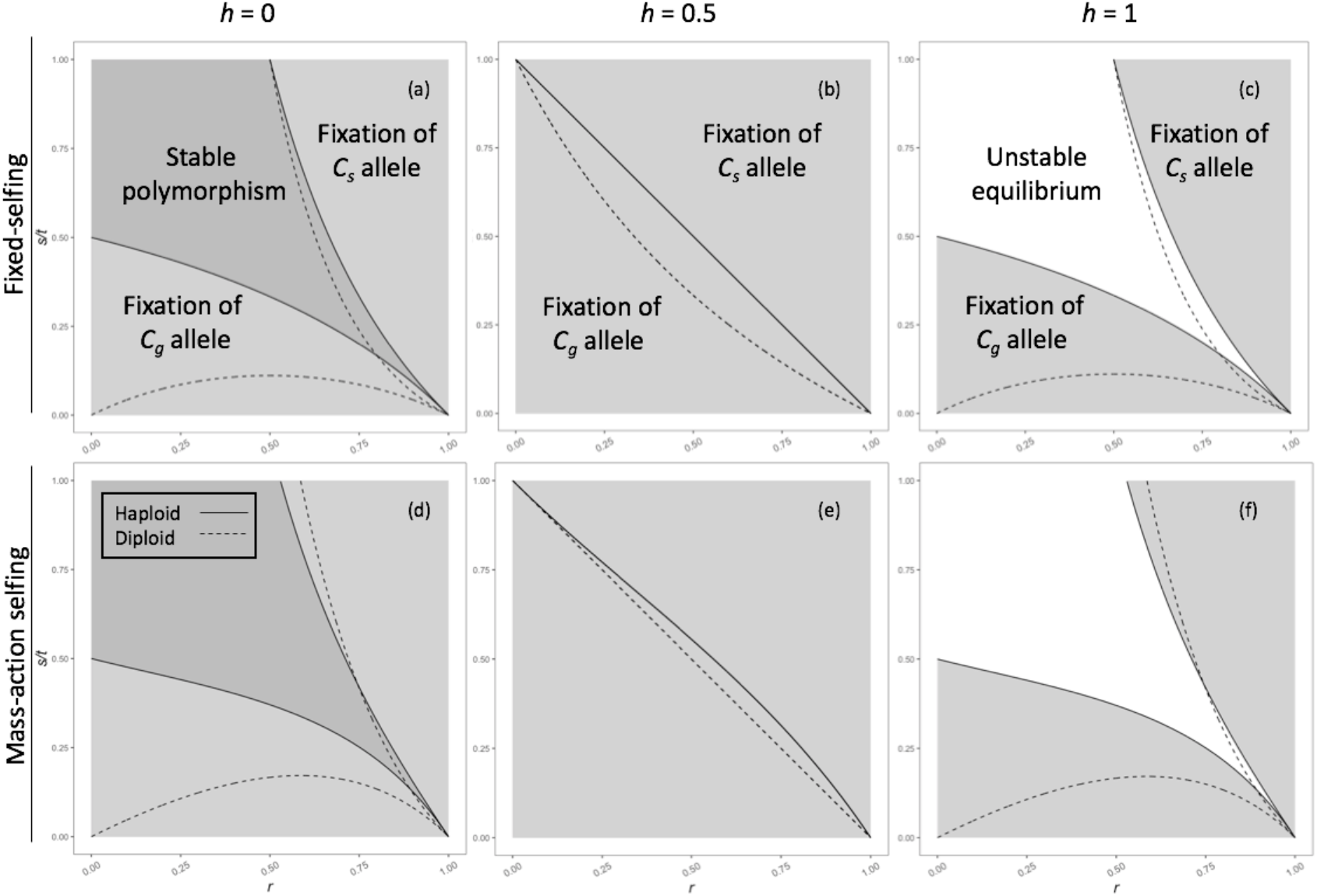
Conditions producing fixation, polymorphism and unstable equilibrium of *C_g_* and *C_s_* alleles under variable diploid selection *s* and dominance *h* for both fixed and mass-action selfing. Analytical solutions producing these conditions do not assume weak selection. Solid lines indicate boundaries under fixed-selfing; dashed lines indicate those for mass-action selfing. Potential for polymorphism diminishes as the dominance of the *C_g_* allele increases, eventually being replaced by unstable equilibrium when *h* > 0.5. As the diploid selection coefficient *s* increases relative to *t*, maintenance of the *C_g_* allele requires either lower dominance *h* or less allocation of reproductive structures to selfing, given by *r*. Compared to fixed-selfing, mass-action selfing conditions permitting polymorphism versus unstable equilibrium are qualitatively similar but quantitatively more limited. The placement of mass-action lines to the right of those for fixed selfing are consistent with individual stigmas never receiving solely self-pollen except when *r* = 1, which increases the opportunity for *C_g_* to exercise its competitive superiority.

Additive fitness effects (*h* = 0.5) largely prohibit polymorphism (Figures 1b, 2). When *h* = 0.5, the critical *r* values for invasion (Eq. 7a-b) reduce to approximately 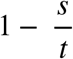 for both alleles.

As *h* increases, the relative fitness of heterozygotes decreases. It then follows that the initial advantage enjoyed by both alleles when rare also diminishes. To invade, *C_g_* requires low *r* values to win enough contests to offset lower heterozygote fitness. *C_s_* will require higher *r* values to avoid competitive losses and produce enough high-fitness *X*_2_ homozygotes to offset lower fitness of heterozygotes.

When the diploid fitness disadvantage of the *C_g_* allele is dominant (*h* = 1), there is a range of selfing rates that prohibit either allele from invading (Figures 1c, 2). For a given value of s, the maximum selfing rate allowing *C_g_* invasion is lowered because its adverse effect on sporophyte fitness is fully expressed by heterozygotes. Oppositely, the critical *r* for *C_s_* invasion increases. When rare, the positive diploid fitness effects of the *C_s_* allele are concealed in heterozygotes. Only very high selfing rates can produce *C_s_* homozygotes quickly enough to provide sufficient refuge from competition, allowing *C_s_* to spread.

### Mass-action selfing

Considering mass-action selfing, the maximum and minimum *r* values allowing invasion of the *C_g_* and *C_s_* alleles can be solved analytically. However, even under weak selection the expressions are complex and difficult to interpret biologically; these are presented in Appendix F and are visualised in Figure (1d-f).

While the expressions for critical *r* values are complex, similarly derived expressions for critical diploid selection *s* values can be used to understand how selfing rate affects invasion success. Expressions for maximum and minimum *s* values permitting invasion of *C_g_* and *C_s_* alleles respectively are given below:

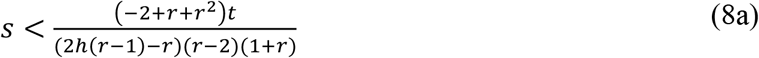

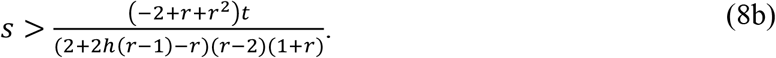

Taking the partial derivative of Eqs. 8a and 8b with respect to *r* lends insight into how these critical *s* values change in response to selfing. (For a discussion of these partial derivative expressions, see Appendix G). These show an increase in selfing rate always decreases the maximum and minimum *s* values permitting invasion of *C_g_* and *C_s_* alleles respectively (Figure 1d-f). It then follows that increased selfing contracts the parameter space allowing invasion of an allele conferring a benefit to pollen competitiveness at the expense of diploid fitness.

### Comparing invasion conditions for fixed and mass-action selfing

Invasion conditions for the two selfing modes were qualitatively similar, with modest quantitative differences. We compared sensitivity of the minimum and maximum *s* values permitting invasion of *C_s_* and *C_g_* alleles to increases in *r* for both systems. (Partial derivative expressions for fixed selfing are also provided in Appendix G). For a given set of *h, s* and *t* values, *r* values allowing *C_g_* and *C_s_* invasion are slightly lower under mass-action selfing relative to those of fixed-selfing (Figure 2), narrowing the permissible parameter space. Thus, under mass-action selfing, the *C_g_* allele can persist at higher selfing rates. Additionally, the range of *r* values permitting polymorphism (when *h* < 0.5) or unstable equilibrium (when *h* > 0.5) contracts under mass-action selfing, which is consistent with narrowed parameter space regions for polymorphism and unstable equilibrium under mass-action selfing observed in Figure 2.

Under fixed selfing, male diploid fitness (pollen quantity) is moot since all flowers of each genotype are assumed to produce enough pollen to fertilise their ovules. Under mass-action selfing, the quantity of pollen produced by an individual affects its realized fertilisation success through self-pollen, as each stigma can receive a mixture of *C_g_*- and *C_s_*-bearing pollen, restricting the competition refuge for *C_s_*. While each *C_g_*-bearing grain has a greater chance of success, this advantage is partially countered by lower pollen production in *C_g_* homozygotes. The different impacts on male function shift the invasion boundary upward under mass-action selfing. For example, when *h* = 0, *s* = 0.01 and *t* = 0.04 (visualised as *s/t* = 0.25 on Figure 1), the maximum selfing rate allowing the *C_g_* allele to invade increases from roughly *r* = 0.8 to *r* = 0.84. Likewise, the minimum selfing rate allowing *C_s_* to invade increases from *r* = 0.67 to *r* = 0.75.

### Fixation and protected polymorphism

Having established invasion conditions for the *C_g_* and *C_s_* alleles, this section explores their equilibrium frequencies. We first present analytical solutions to the recursion equations (Eqns. 3a-c, 5a-c) assuming weak selection (0 < *s, t* ≪ 1). We then highlight the difference in outcomes when pollen competitiveness is under gametophyte genetic control, as Haldane speculated, versus sporophyte control (seen in Jordan and Connallon 2014). Under fixed selfing, the equilibrium value 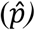 of the *C_g_* allele is given as:

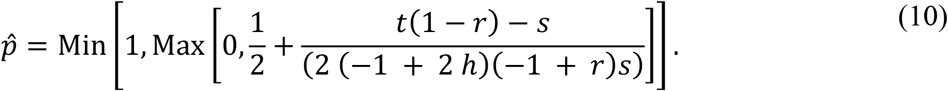

When *C_g_* is recessive (*h* < 0.5) and mating is mixed (0 < *r* < 1), the denominator of the far-right term is always positive. Whether 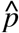 falls above or below 0.5 then depends on the sign of the numerator. If *r* < 1 − *s/t*, then 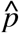 will fall above 0.5 but fall below if the inverse is true. Simply, the higher the selfing rate (*r*) the higher the diploid cost (*s*) needed to prevent *C_g_* fixation, which is implied by a 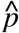 value < 0.5. When *C_g_* is dominant (*h* > 0.5), the above expression gives the repelling equilibrium frequency (above 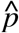 the *C_g_* allele goes to fixation and below is eliminated). When *h* > 0.5 and mating is mixed, the denominator of the far-right term is always negative. Thus, the repelling 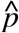 value falls above 0.5 when the numerator is negative and below when positive. Looking at the effect of selfing rate on 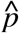, the threshold 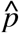 allowing *C_g_* invasion decreases as *r* decreases, as the numerator will become increasingly positive. When *h* = 0.5, the denominator of the far-right terms goes to zero, putting the whole term to infinity. Threshold selfing rates permitting or preventing *C_g_* invasion are thus given by the sign of infinity, which is determined by if *r* is greater than or less than 1 − *s/t* respectively.

Under mass-action selfing, the equilibrium frequency of the *C_g_* allele is:

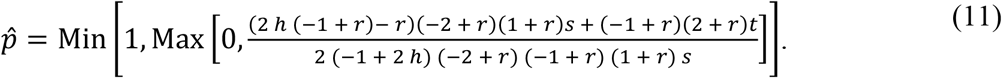

When *h* < 0.5, Eqs. 10 and 11 provide the equilibrium frequency of the *C_g_* allele. When *h* > 0.5, these expressions provide the ‘repelling’ allele frequency. Figure 3 plots the equilibrium frequency of *C_g_* as a function of selfing rate, *r*, under relatively weak selection. (For a comparison of analytical results to numerically derived frequencies under strong selection, see Appendix E).

**Figure 3.**
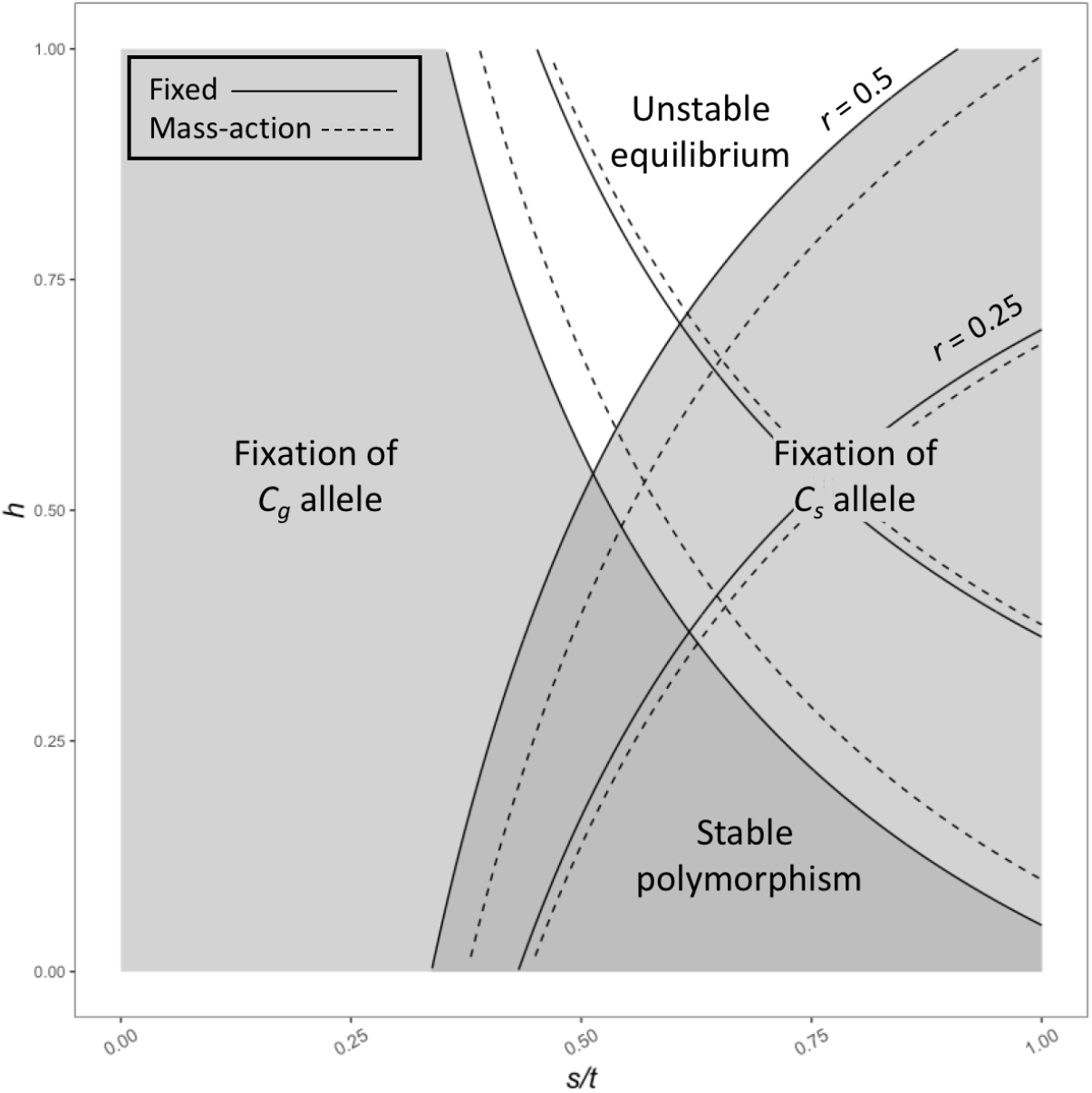
Analytical results of equilibrium frequency of the *C_g_* allele across selfing rates 0 < *r* < 1 under gametophytic (haploid) and sporophytic (diploid) genetic control. Analytical solutions were derived assuming weak selection. Solid lines indicate equilibrium allele frequencies, while dashed lines indicated the analytical repelling frequency (i.e., the starting *p* frequency below which*p* will be lost and above which it will fix). (a-c) Under the fixed-selfing model, the window of selfing rates contracts as dominance increases. (d-f) Under the mass-action model, *C_s_* pollen is never entirely protected from competition with *C_g_* pollen as individual stigmas receive a mixture of both self and outcross pollen for intermediate selfing rates. Thus, maximum and minimum selfing rates allowing fixation or maintenance of *C_g_* and *C_s_* alleles respectively are right-shifted compared to under fixed selfing, reflecting the greater opportunity for the *C_g_* allele to exercise its advantage in pollen competition with the *C_s_* allele. Notably, for both selfing modes, the range of selfing rates at which the *C_g_* allele can invade contracts under diploid genetic control relative to haploid control.

When the negative diploid effect of *C_g_* is dominant (*h* = 1), high selfing prevents *C_g_* invasion when rare, while low selfing prevents *C_s_* invasion. Under intermediate selfing rates, neither allele can increase when rare (Figure 3c, f). Following the appearance of the *C_g_* allele in a population by mutation, its initial frequency will almost inevitably fall below the repelling frequencies predicted by Eqs. 10 or 11 and thus it will be eliminated.

Comparison of Eqs. 10 and 11 reveals that when 0 < *h* < 0.5, the equilibrium frequency 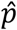 is higher under mass-action selfing relative to fixed selfing; however, when *h* > 0.5, the inverse is true (for derivation, see online supplemental materials). As suggested by invasion conditions, under mass-action selfing, minimum and maximum selfing rates allowing invasion of *C_g_* and *C_s_* alleles respectively increase for given *h, s* and *t* values (Figure 3d-f). Additionally, parameter space allowing both polymorphism (Figures 3d) and unstable equilibrium (Figures 3f) contracts under mass-action selfing relative to fixed (Figures 3a, 3c).

As is visualised in Figures 1 and 3, invasion conditions and equilibrium allele frequencies under both selfing modes assuming haploid control of pollen competitiveness are qualitatively similar to those derived using models provided in Jordan and Connallon (2014), where competitiveness is under paternal genotype control (for derivation of analytical results, see online supplemental materials; for models and discussion of analytical solutions, see Appendix B). They also found selfing and dominance restrict potential for stable polymorphism and ultimately diminish selection through male function. Our results show, however, that gametophytic control substantially expands the conditions favouring an allele promoting pollen competitiveness. With gametophyte gene expression, pollen phenotype is more strongly correlated with haplotype. Heterozygotes always produce a 50:50 mix of phenotypically and genetically ‘strong’ and ‘weak’ pollen competitors. Under sporophytic control, heterozygotes can produce phenotypically all strong (*h* = 0), all weak (*h* = 1), or all intermediate (*h* = 0.5) competitors, despite always producing a 50:50 genetic mix. Further, under gametophytic control, variance in competitive ability between pollen carrying opposite alleles is constant. Under paternal control, variance in competitive ability changes with genotype frequency: increasing heterozygote frequency decreases the difference in mean competitiveness of the two pollen haplotypes. Consequentially, *C_g_* invasion occupies a markedly smaller parameter space when under diploid genetic control (Figure 1). By extension, we see that the *C_g_* allele reaches a lower equilibrium frequency, *i.e*., invades with more difficulty, for a given selfing rate when it is expressed in the sporophyte rather than the gametophyte (Figure 3).

Returning to Haldane’s conjecture that plants are at the mercy of their pollen grains, how strong must sporophyte selection against a pollen-competition allele be to prevent its fixation? The fate of the *C_g_* allele depends on selection strength during the two life stages, which can be considered as a ratio of diploid and haploid selection coefficients, *s/t*. At a given *s/t* ratio, increasing the selfing rate will restrict opportunities for *C_g_* pollen to exercise its superiority, muting the selective advantage. Against this, at any given ratio, Mendelian dominance alters how often the diploid cost is paid. If *C_g_* is recessive (*h* = 0), heterozygotes pay no diploid cost. If dominant (*h* = 1), it masks the *C_s_* benefit and heterozygotes pay full cost; however, since selfing decreases heterozygote frequency, the dominance issue becomes moot. These conditions produce the general outcome that *s/t* ratios allowing *C_g_* to invade, and to be maintained in polymorphism, tend to decrease as both *r* and *s* increase (Figure 4).

**Figure 4.**
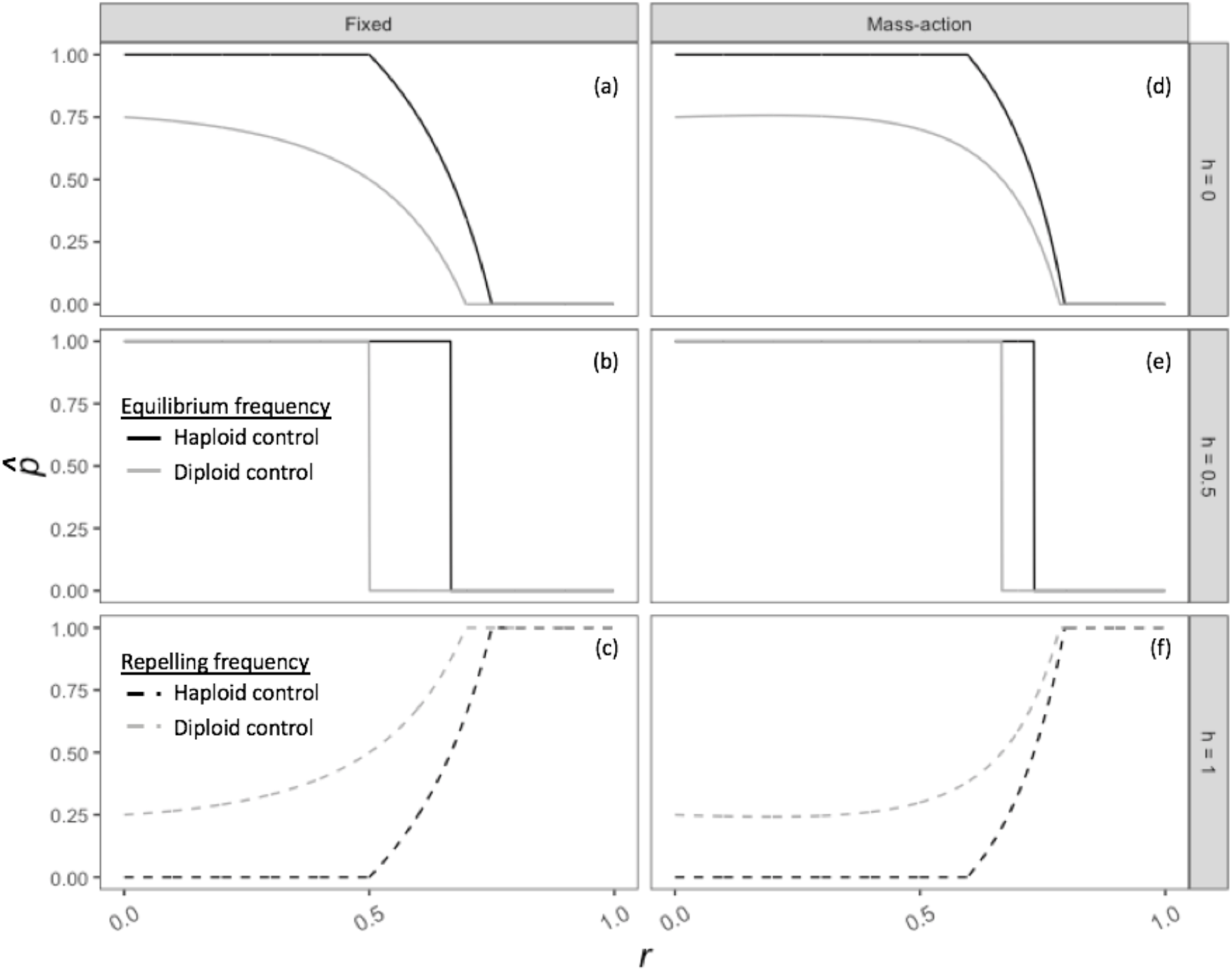
Analytical results for equilibrium frequency of the *C_g_* allele across a range of *s/t*, i.e., ratios of diploid and haplod fitness values. Series specify selfing rates *r* ranging between 0 and 1. Under both (a-c) fixed and (d-f) mass-action selfing models, increases in dominance of the *C_g_* allele sharply decrease the maintenance of polymorphism, particularly at lower selfing rates. Noticably, for a given combination of selfing rate *r* and dominance *h*, the *C_g_* allele persists at higher ratios of *s* and *t* under mass-action selfing relative to fixed-selfing, reflecting the greater opportunity for *C_g_*-bearing pollen to compete *C_s_*-bearing pollen under mass-action selfing. For both selfing models, regardless of dominance, the *C_g_* allele cannot invade under complete selfing, i.e., *r* = 1.

**Figure 5.**
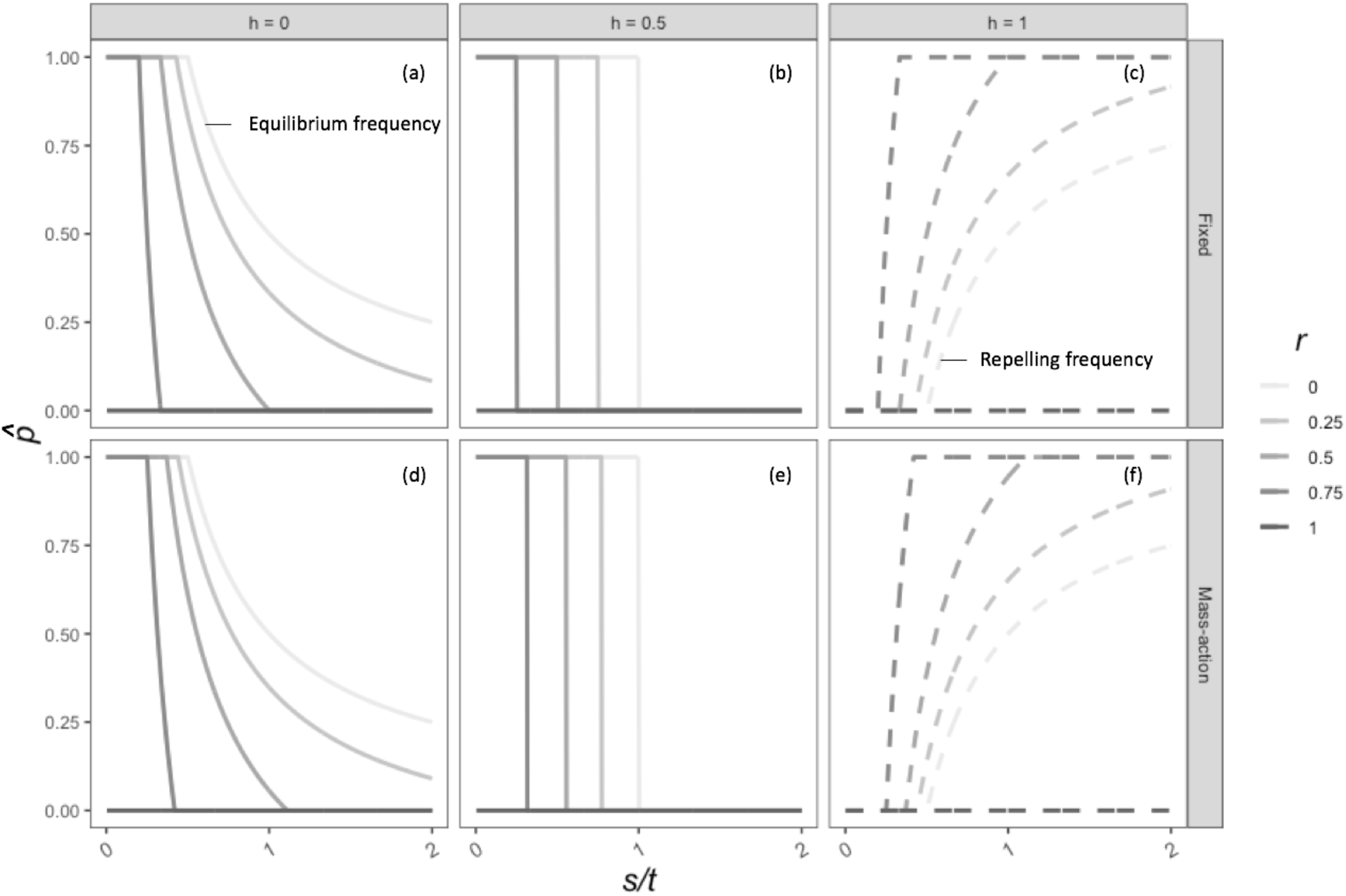

## DISCUSSION

Classical theory, which assumes random mating, predicts an allele conferring an advantage in pollen competitiveness fixes unless a trade-off with sporophyte fitness sufficiently counters this haploid advantage. Our models predict that even moderate selfing can prevent the spread of a pollen-expressed allele that confers competitive superiority (Table 1). Increased selfing increases assortative competition among pollen, restricting opportunity for a competitive allele to exercise its superiority. When homozygotes self completely, they deposit only one pollen haplotype per stigma, so by necessity that haplotype cannot be out-competed by another. Alternate haplotypes compete only on heterozygotes. Further, selfing depresses heterozygote frequency such that the proportion of the pollen pool engaged in competitive contests falls in proportion to the selfing rate.

**TABLE 1.** Summary of model outcomes for mutation to an allele that increases pollen (haploid gametophyte) competitive ability but reduces survivorship or fertility in the (diploid) sporophyte.

| EXPRESSION OF *C<sub>g</sub>* IN DIPLOID SPOROPHYTE | MATING SYSTEM |  |  |
| --- | --- | --- | --- |
|  | Obligate Outcrossing (*r* = 0)<br>*Ample opportunities to exercise pollen competitive superiority* | Mixed (0 < *r* < 1)<br>*Fewer opportunities to exercise pollen competitive superiority* | Obligate selfing (*r* = 1)<br>*No opportunities to exercise pollen competitive superiority* |
| **Recessive**<br>Fitness cost paid only by homozygote | *C<sub>g</sub>* goes to fixation if sporophyte cost is less than ½ the pollen competitive benefit, otherwise reaches stable polymorphism– *conditions for stable polymorphism broad.* | *C<sub>g</sub>* goes to fixation at increasingly lower sporophyte costs as selfing rate increases; upper limit for cost to permit polymorphism is increasingly reduced at higher selfing rates– *conditions for stable polymorphism narrowed.* | *C<sub>g</sub>* unable to invade. |
| **Dominant**<br>Fitness cost paid by *both* homozygote and heterozygote | *C<sub>g</sub>* goes to fixation if sporophyte cost is less than ½ the pollen competitive benefit, but eliminated if costs are higher – *no stable polymorphism.* | *C<sub>g</sub>* goes to fixation at increasingly lower sporophyte costs as selfing rate increases, but *eliminated* if costs are any higher – *no stable polymorphism.* | *C<sub>g</sub>* unable to invade. |

The dominance relationships between the competitive and non-competitive alleles in the diploid stage also influence invasion. When the sporophyte fitness cost of the *C_g_* allele is recessive, it is shielded from negative selection during the diploid phase. Complete dominance fully exposes it to negative selection. Thus, the conditions for invasion narrow with increases in the dominance coefficient (Table 1, Figures 1–4).

Increased dominance also narrows the condition for stable polymorphism (Table 1, Figures 1–4). When costs are recessive, heterozygotes gain success as fathers because 50% of their pollen is competitively superior and they pay no cost. This can lead to overall heterozygote superiority. Under random mating, this over-dominance in total fitness can maintain stable polymorphism; however, selfing diminishes the impact of over-dominance by eroding heterozygote frequency: heterozygosity lost to segregation is not replenished by matings between opposite homozygotes, and so the competitive allele is eventually carried mostly by the homozygote, who pay full cost. If the competitive allele is dominant, stable polymorphism is precluded. In this case, the heterozygotes still get half the paternal benefit but pay full cost (Table 1, Figures 1–4), making over-dominance for total fitness unachievable.

By comparing invasion conditions and equilibrium allele frequencies under haploid and diploid control (as seen in Jordan and Connallon 2014) of pollen ability (Figures 1, 3), we demonstrate that the form of genetic control overseeing competitive ability has important implications for the fate of an allele conferring an advantage to pollen ability. The stronger haplotype-phenotype correlation in our model is important during the early stages of invasion, when pollen carrying the competitive allele is produced only by heterozygotes: it assures that the pollen grains carrying the competitive allele are indeed competitive. This clear genetic signal expands conditions for a costly allele to invade and spread; stronger selfing is required to counter the advantage.

Our fixed and mass-action selfing models are caricatures representing two extremes in pollination systems, but real systems can be intermediate. In *Impatiens palida*, cleistogamous flowers always self whereas there is a small incidence of geitenogamous selfing in chastogamous flowers (Schemske 1978). Gradations in selfing prior to bud opening or delayed self-pollination when pollinator visitation is low (Loyd 1979) can generate variation in the proportion of self-pollen involved with competitive contests with outcross. However, we have shown strong qualitative agreement between the outcomes of the two extreme models, both leading to the general prediction that selection favouring increased pollen competitive ability is weakened by selfing.

Genomic analysis of exclusively pollen-expressed loci expressed, where diploid costs are necessarily zero, bears out the prediction of relaxed selection under self-pollination. In the self-incompatible *Capsella grandiflora*, evidence points to stronger purifying selection at sites expressed solely in the pollen tubes than at sites expressed exclusively in the seedling (Arukumar et al., 2011). In the highly selfing *Arabidopsis thaliana*, by contrast, purifying selection on pollen-exclusive genes is weaker than on sporophyte-exclusive genes, as evidenced by a higher frequency of premature stop codons in the former (Harrison et al. 2015). Greater scrutiny of loci expressed during both stages are needed to clarify the general importance of negative pleiotropic constraints on the evolution of gametophyte and sporophyte traits.

## Figure captions

**Figure 1** Conditions producing fixation, polymorphism and unstable equilibrium (i.e., fixation is dependent on initial frequencies) of two alleles *C_g_* and *C_s_* at an autosomal gene under antagonistic pleiotropy between pollen competitive ability and diploid fitness. Solid lines delineate conditions under gametophytic genetic control and dashed lines under sporophytic genetic control. (a-c) Analytical results for the fixed-selfing model assuming weak selection. Potential for polymorphism diminishes as the dominance of the *C_g_* allele increases, eventually being replaced by unstable equilibrium when *h* > 0.5. As selfing rate (*i.e*., proportion pollen dedicated to selfing) increases, polymorphism requires greater diploid selection *s* against the *C_g_* allele to offset its greater pollen competitive ability *t*. (d-f) Analytical results assuming weak selection for the mass-action model, where *r* is the proportion pollen dedicated to selfing and stigmas receive a mixture of self- and outcross pollen. Conditions allowing invasion of the *C_g_* allele are notably greater than under fixed selfing, as individual stigmas never receive entirely self-pollen except when *r* = 1, thus increasing opportunity for competition between *C_g−_* and *C_s−_* bearing pollen. For both fixed and mass-action selfing, relative to haploid control, diploid control expands conditions under which the *C_s_* allele can be maintained while contracting those under which the *C_g_* allele can be maintained.

## Appendices

## Appendix A

### Implications of sex-specific dominance

Given we assume diploid selection acts either through lower flower production or reduced survival, we also assume a common dominance coefficient *h* for female and male function (i.e., quantities of ovule and pollen produced). However, it is possible to consider the effects of different dominance coefficients for male and female function, given as *h_m_* and *h_f_*. In such a case, absolute fitness via female function for *C_g_C_s_* individuals is *W_1f_* = 1−*h_f_s* and that via male function is *W_1m_* = 1−*h_m_s*. Absolute fitness terms for homozygote individuals remain unchanged from the main text. Using these sex-specific dominance coefficients, we can derive new expressions for equilibrium frequency of the *C_g_* allele under weak selection for both fixed (Eq. A1) and mass-action (Eq. A2) selfing:

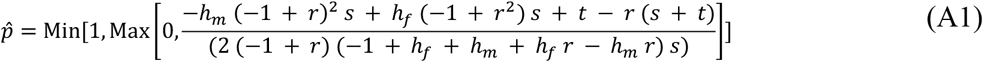

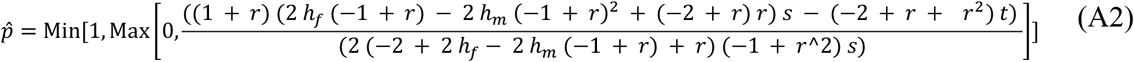

To understand the relative impacts of changing *h_f_* and *h_m_* on 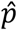, we can take the partial derivative of the above equations first with respect to *h_f_* and then with respect to *h_m_*. For fixed selfing, these partial derivatives are as follows:

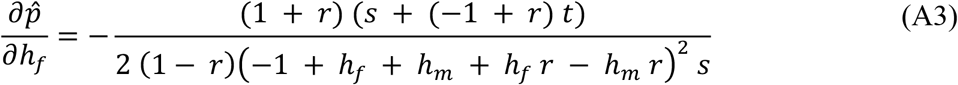

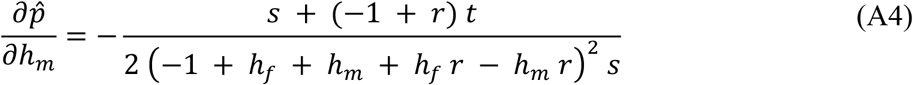

With simple arranging, it can be shown that

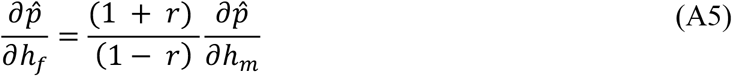

In a mixed mating system where 0 < *r* < 1, the far-left term of the right-hand side of Eq. A5 is always greater than 1 and increasing as selfing rate increases, meaning an increase in *h_f_* always yields a greater change in 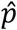 than a comparable increase in *h_m_*. This is expected, as under fixed selfing, the cost to male diploid fitness is only paid through outcrossing, as we assume all plants produce enough pollen to self-fertilise. Conversely, diploid fitness effects on female function are recognised both in selfing and outcrossing. Thus, the effects of changing female and male dominance terms are only equivalent under complete outcrossing (*r* = 0) and Eq. A5 reduces to 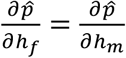.

We can also compare the effect of increasing *h_f_* relative to a shared dominance coefficient *h*. Returning to Eq. A3, it is evident that the denominator is always positive, while the sign of the numerator changes from positive to negative when *r* switches from less than 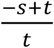 to more than 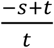. This particular value, 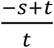, gives the selfing rate *r* at which 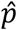 is 0.5. Looking specifically at the effects of dominance, the magnitude of the slope of 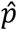 in response to *r* can be controlled by adjusting *h_f_* in the denominator. Increasing the value of *h_f_* increases the magnitude of the denominator, thus yielding a more positive slope when 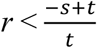 and a more negative slope when 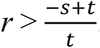. This effect is visualised in Figure A1. Compared to *h_f_* = *h_m_* = 0, increasing *h_f_* to 0.25 allows *p* to remain fixed at higher selfing rates but also to be lost more quickly, thus contracting the space for polymorphism. An identical effect, but for space allowing unstable equilibrium, is observed when all dominance coefficients are above 0.5 (Fig. A2).

**Figure A.1.**
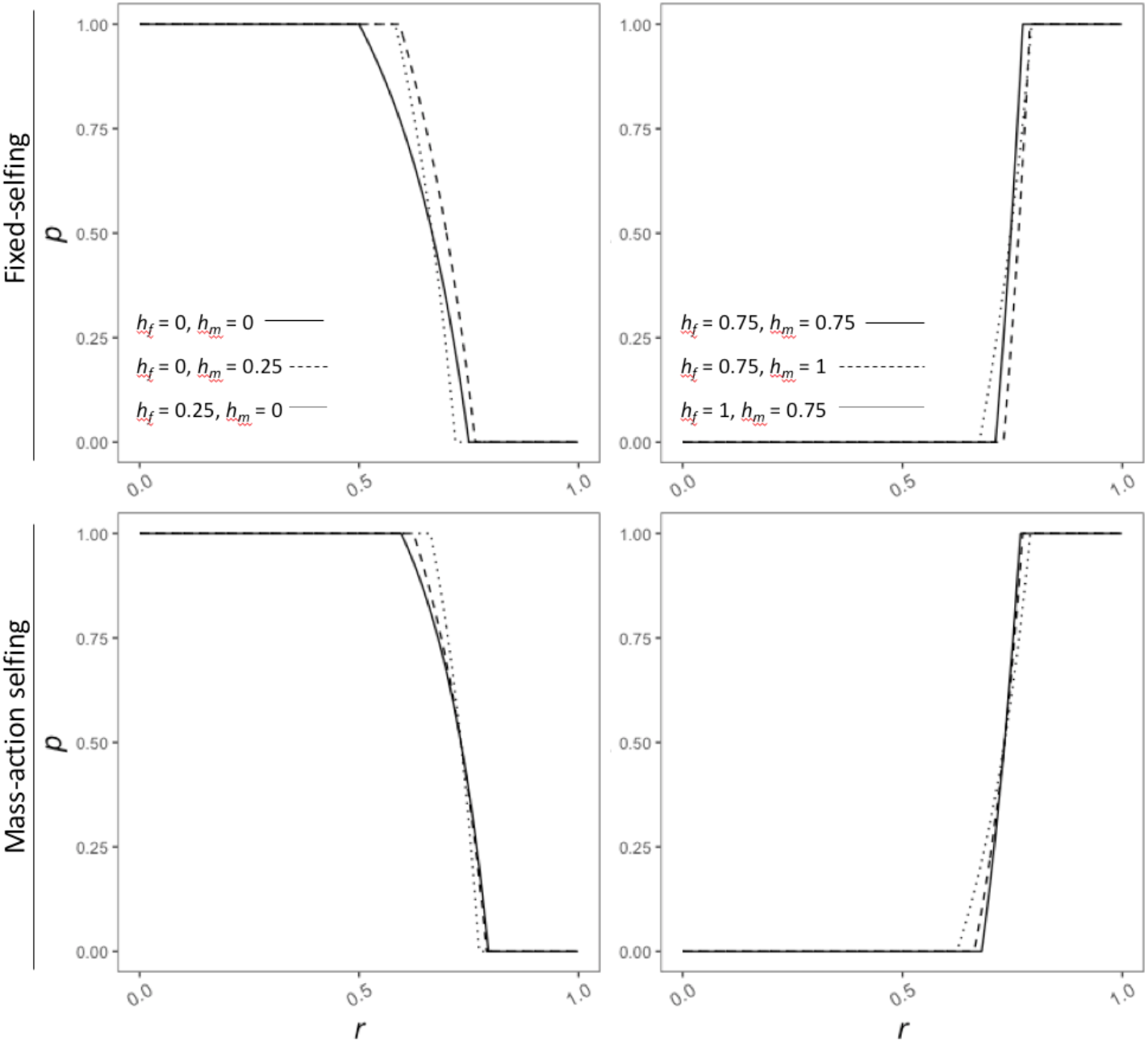

When *h_f_* increases and the *C_s_* allele is rare, the *C_s_* allele confers a lesser advantage to female diploid fitness as this advantage is increasingly masked by the *C_g_* allele in heterozygotes (in which it is predominantly found when rare). As an increase in *h_f_* means the *C_s_* allele enjoys less of a diploid fitness benefit over the *C_g_* allele when found in heterozygotes, the *C_s_* allele becomes increasingly dependent on the production of homozygotes to facilitate its invasion.

Thus, as *h_f_* increases, *C_g_* remains fixed up to higher values of *r*. A similar logic follows for invasion of the *C_g_* allele. When *h_f_* increases, the *C_g_* allele also suffers in the heterozygote. It thus requires lower selfing rates so that it can exercise its pollen superiority competitive over the *C_s_* allele under outcrossing.

When *h_m_* increases, qualitatively similar but quantitatively different effects occur. Grouping *h_f_* terms together in the denominators of Eqs. A3 and A4 produces *h_f_* (1+*r*), while grouping *h_m_* terms together produces *h_m_* (1−*r*). This means that an increase in *h_m_* produces a smaller change in the size of the denominator than an equivalent change in *h_f_*. This is expected, as diploid fitness effects through male function are only experienced under outcrossing, unlike female fitness effects, which are experienced both under selfing and outcrossing. Otherwise, the logic of how dominance affects invasion of both *C_g_* and *C_s_* alleles described above for *h_f_* also applies to *h_m_*, but to a lesser magnitude. This difference is visualized in Figure A1, as the lines depicting *h_f_* = 0.25 and *h_m_* = 0.25 both remain fixed for*p* up to higher selfing rates, but also drop (and thus lose *p* entirely) at lower selfing rates, when compared to *h_f_* = *h_m_* = 0.

Turning to mass-action selfing, the partial derivatives of Eq. A2 with respect to *h_f_* (Eq. A6) and *h_m_* (Eq. A7) are as follows:

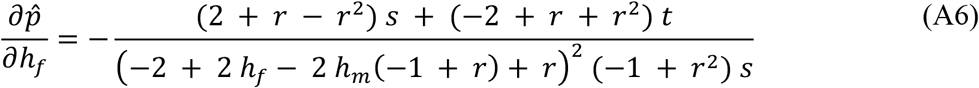

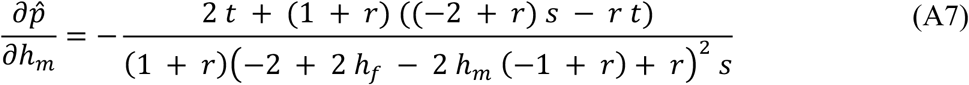

Rearrangement of Eqs. A6 yields:

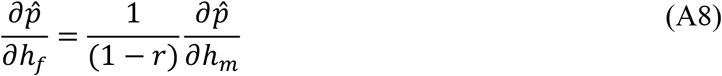

As seen under fixed selfing, an increase to *h_f_* will yield greater change to 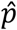 than an equivalent change to *h_m_*. Interestingly, the ratio of these changes is not the same between fixed and mass-action selfing. Relative to a change in *h_m_*, an equivalent increase in *h_f_* under mass-action selfing has a weaker effect on *p* than under fixed selfing, as evidenced by comparing the coefficients of Eqs. A5 and A8, 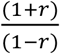 and 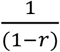 respectively (assuming *r* > 0). This can be understood by considering where diploid selection on male function acts in mass-action selfing. Unlike fixed selfing, costs to pollen production are paid both through both self-pollen and outcross-pollen, as both pollen types compete on the same stigma. Differences in pollen competitive ability offset costs through diploid male fitness, thus allowing an increase in *h_m_* to still have less impact on 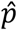 than an increase in *h_f_*. The lesser effect of altering *h_m_* on 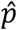 can be seen in Figure A1. Compared to fixed selfing, increasing *h_m_* produces less deviation from baseline series (*i.e*., those with equivalent *h_m_* and *h_f_* values).

## Appendix B

### Models for sporophytic control of pollen competitive ability (Jordan and Connallon 2014)

The relative frequencies of each genotype among the available ovules are given as:

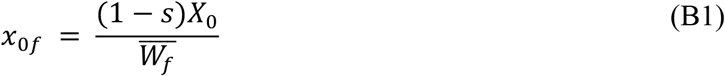

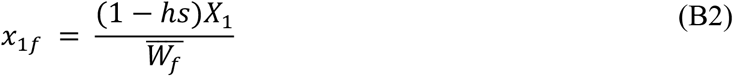

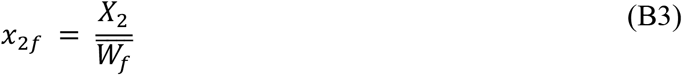

Where *s* is the selection coefficient acting on female function, *h* is the dominance coefficient and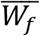 is defined as

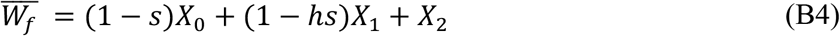

The relative frequencies of each genotype among the pollen pool are given as:

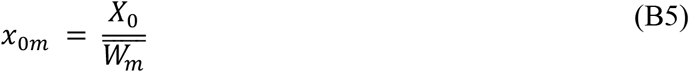

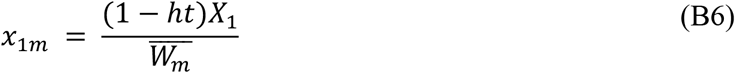

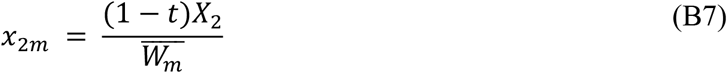

Where *t* is the selection coefficient acting on male function, *h* is the dominance coefficient and 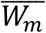 is defined as

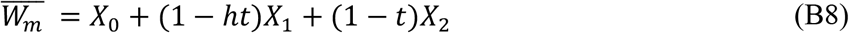

Using these fitness expressions, the genotype recursion equations are:

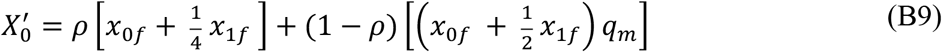

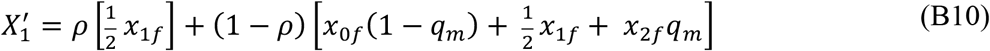

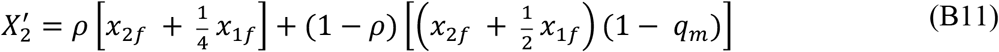

where *q_m_* is the frequency of the *C_g_* allele in the pollen pool:

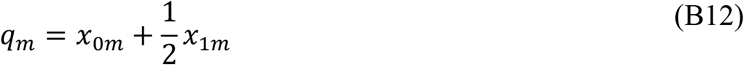

In the case of fixed selfing, selfing rate *ρ* is equivalent to *r* (i.e., proportion pollen allocated to selfing). Under mass-action selfing, *ρ* is frequency-dependent and genotype-specific:

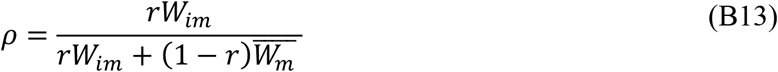

where *W_0m_* = 1, *W_1m_* = (1 − *ht*) and *W_2m_* = 1 − *t* and *r* is the proportion of pollen dedicated to selfing.

Equilibrium allele frequencies for fixed and mass-action selfing assuming sporophytic control were derived using the same methods provided in the main text and are as follows:

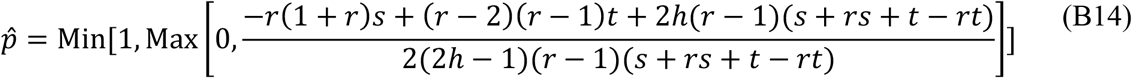

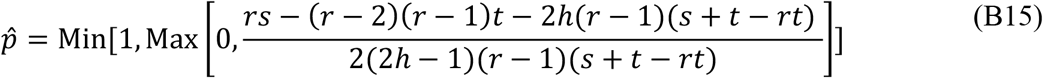

We can understand the conditions under which the *C_g_* allele will perform better under sporophytic control relative to haploid and *vice versa* by taking the difference between Eqs. 10 (main text) and B16 as well as Eqs. 11 (main text) and B15. Under fixed selfing:

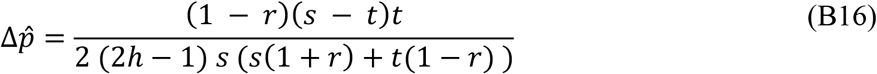

In Eq. B16, when *h* < 0.5, the denominator is always negative. The sign of the numerator is dependent on the size of *s* relative to *t*. When diploid selection *s* is larger than haploid selection *t*, the numerator is positive and thus 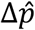 is negative, revealing the *C_g_* allele is more successful under sporophytic than gametophytic genetic control. Inversely, when *s* < *t* (as presented in the main text), 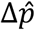 is positive and *C_g_* reaches a higher frequency under gametophytic control.

Under mass-action selfing:

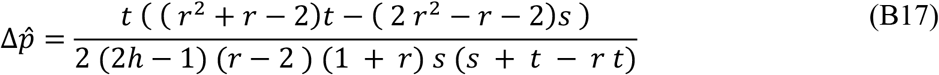

When *h* < 0.5 and 0 ≤ *r* ≤ 1, the denominator of B17 is negative. Thus, the sign of 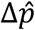 is dependent on the sign of the numerator. Given 0 ≤ *r* ≤ 1, (*r*^2^ + *r* − 2) is always negative and (2 *r*^2^ − *r* − 2) is always positive, meaning the numerator is always negative and 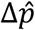 is always positive. In other words, when *h* < 0.5, the *C_g_* allele will always do better under gametophytic control than under sporophytic control. Conversely, when *h* > 0.5, the denominator is positive while the numerator remains unchanged. In such a case, 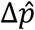 is always negative and the *C_g_* allele reaches a higher equilibrium frequency under sporophytic control. These analytical results are visualised in Figure 3, in which, when *h* = 0, the equilibrium allele frequency 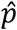 is higher for gametophytic (i.e., haploid) control relative to sporophytic and, when *h* = 1, the repelling allele frequency below which *C_g_* cannot invade is higher under sporophytic control (meaning fixation requires a higher initial allele frequency). This result can be understood easily by considering the haplotype-phenotype correspondence under both gametophytic and sporophytic genetic control. Under gametophytic control, this correspondence is complete (i.e., phenotype reflects haplotype and the *C_g_* allele always confers a benefit to pollen). Under sporophytic control, this correspondence is a function of the paternal genotype (homozygote or heterozygote) and the dominance coefficient h, and thus, the *C_g_* allele will not always confer its maximum advantage to pollen ability, weakening its invasion ability.

Likewise, invasion conditions for fixed and mass-action selfing assuming sporophytic control were derived using the methods found in the main text. Under fixed selfing, maximum and minimum selfing rates permitting invasion of *C_g_* and *C_s_* alleles respectively are given as:

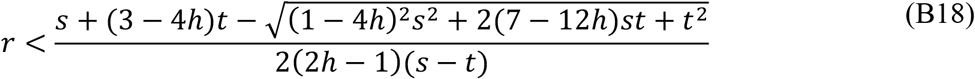

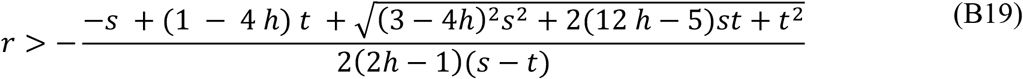

The corresponding expressions for mass-action selfing are:

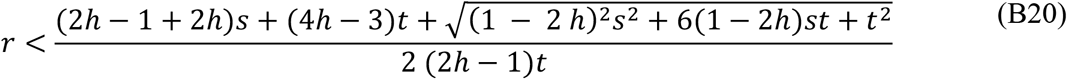

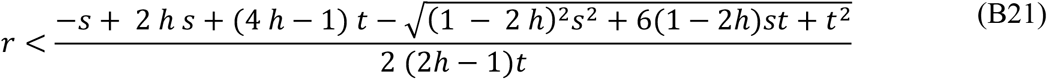

## Appendix C

### Production of heterozygote offspring via selfing with variable pollen competitiveness

With a simple Punnet square, it can be demonstrated that even with differential competitive ability between pollen haplotypes, selfing of heterozygotes produces ½ heterozygote offspring:

**Table C 1.**

| | **$C_g$ pollen**<br>$\frac{1}{2-t}$ fertilisation success<br>given 50% allele frequency | **$C_s$ pollen**<br>$\frac{1-t}{2-t}$ fertilisation success<br>given 50% allele frequency |
| --- | --- | --- |
| **$C_g$ ovule**<br>0.5 allele frequency | $\frac{1}{2(2-t)} C_g C_g$ | $\frac{1-t}{2(2-t)} C_g C_s$ |
| **$C_s$ ovule**<br>0.5 allele frequency | $\frac{1}{2(2-t)} C_s C_g$ | $\frac{1-t}{2(2-t)} C_s C_s$ |

Together, the frequency of heterozygote offspring produced via heterozygote selfing is 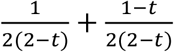, which reduces to ½.

## Appendix D

### Analytical results for mass-action selfing with pollen discounting

Pollen discounting, i.e., loss of pollen for outcrossing during transport, can be incorporated by inclusion of an additional term, *d*, which gives the proportion of pollen lost and is identical to the pollen discounting term included in Jordan and Connallon (2014). Equations 5a-c are thus modified to be:

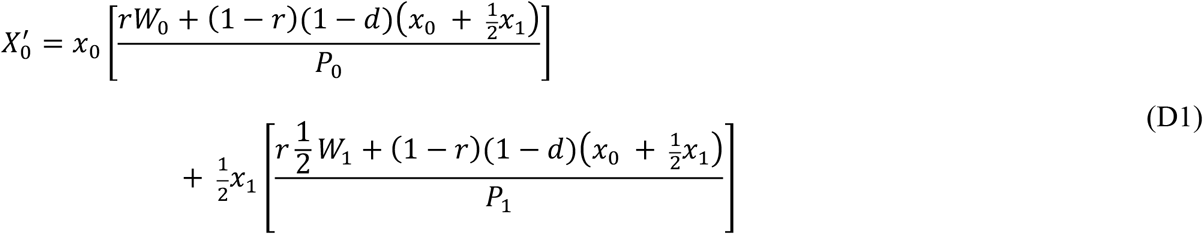

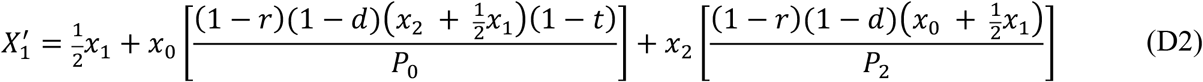

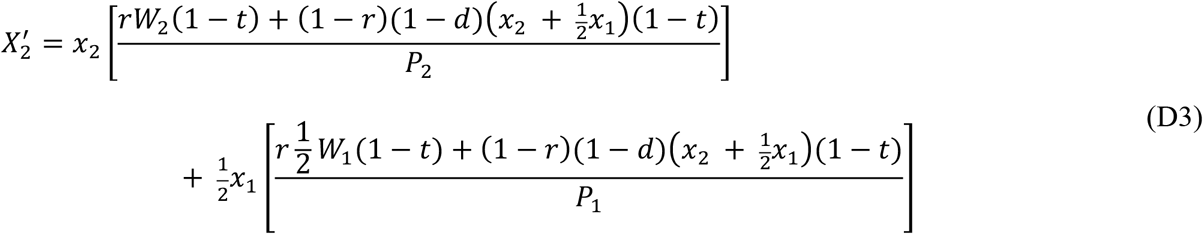

Similarly, Eqs. 6a-c are also modified:

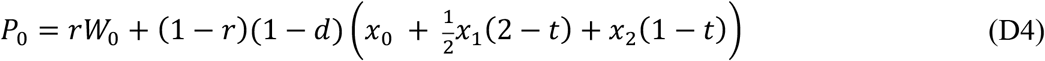

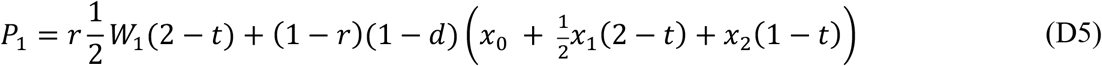

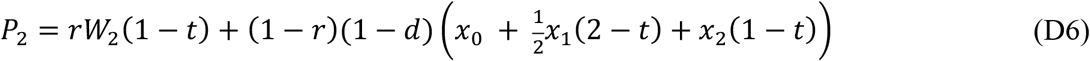

Using Eqs. D1–6, and assuming weak selection, we can derive the equilibrium frequency of the *C_g_* allele. For the full expression, see the corresponding online supplemental materials. To understand the effects of increasing *d* on 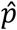, we take the partial derivative of the 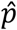 expression with respect to *d*:

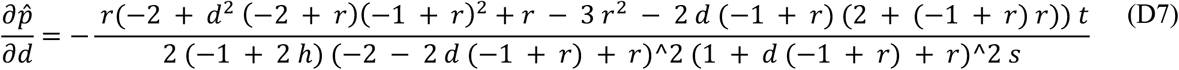

To get a better handle on the above expression, we can isolate and rearrange the terms in which *d* appears to:

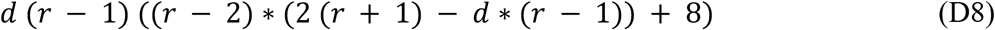

The term *d*(*r*−1) is always negative (when 0< *r* <1), so how *d* affects the equilibrium allele frequency is dependent on the sign of the following term:

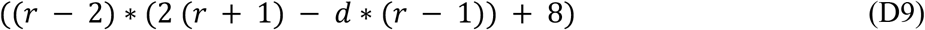

With further rearrangement, it can be shown that:

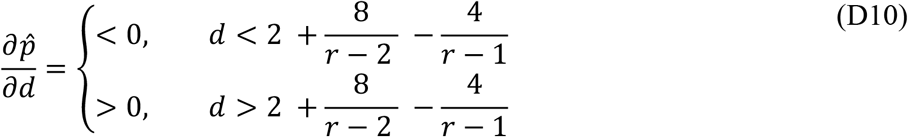

As both *d* and *r* are constrained to between 0 and 1, 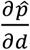 is always positive and thus an increase in *d* always lowers the equilibrium frequency of the *C_g_* allele. This is not surprising as inclusion of a pollen discounting term *d* > 0 should decrease the relative proportion of outcross-versus self-pollen that a stigma receives, therefore decreasing the realised competitive advantage of *C_g_* pollen, under the mass-action selfing model.

In other words, under mass-action selfing, intermediate *r* values indicate a stigma receives both self and outcross pollen. Successful invasion and fixation of either the *C_g_* or *C_s_* allele depends on the ratio of self to outcross pollen produced, with higher values of *r* favouring the *C_s_* allele. Pollen discounting essentially alters this ratio, allowing the *C_s_* allele to invade and fix at lower values of *r*, which thus implies limited invasion of the *C_g_* allele.

Figure D1 displays this effect of pollen discounting on the success of the *C_g_* allele using *t* = 0.1, *s* = 0.05 and *d* values of 0, 0.2 and 0.5. Indeed, as *d* is increased, the equilibrium frequency of the *C_g_* allele becomes increasingly left-shifted, implying a decreased range of selfing rates at which it can successfully invade.

**Figure D 1.**
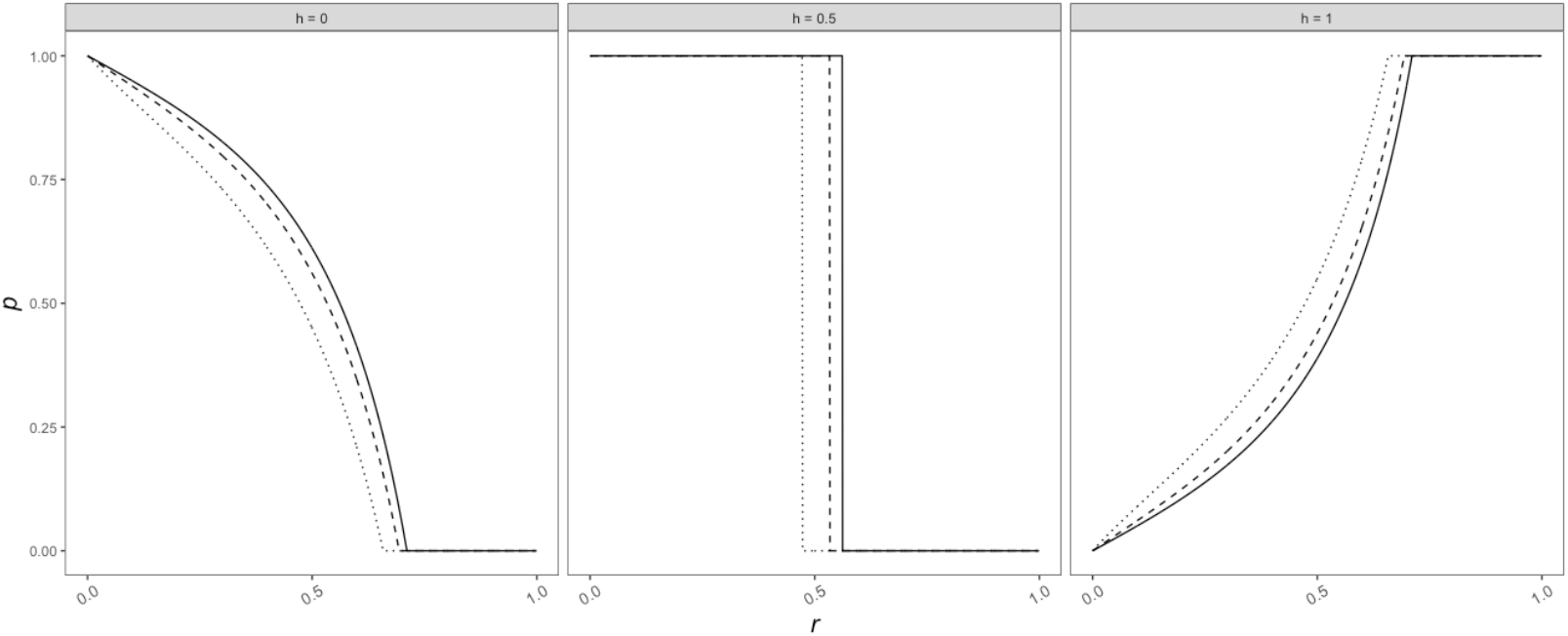

## Appendix E

### Numerical validation and consideration of strong selection

Equilibrium and repelling allele frequencies presented in the main text are entirely analytical, i.e., derived using Equations 10 and 11. Confirmation of these analytically derived frequencies was done numerically. All numerical results were produced using RStudio Version 1.1.353, in which we generated and recorded genotype and allele frequencies over generations for different combinations of selfing, diploid fitness and pollen competitiveness. For invasion from *X*_0_ and *X*_2_ boundaries, initial frequency of the invading allele was always set to 0.0001, and initial genotype frequencies were at Hardy-Weinberg equilibrium. Equilibrium allele frequencies presented in Figures E1 were evaluated at selfing rates *r* ranging from 0 to 1 in increments of 0.01. Long-term allele frequencies were taken from the 50,000^th^ generation and verified by a change in allele frequency, i.e., Δ*p*, of less than 10^−6^. We consider these long-term allele frequencies to be equilibrium frequencies, as we assume that, with such significantly small changes in allele frequencies, drift will dominate, and these long-term frequencies are good approximations of true equilibrium frequencies, where Δ*p* = 0. As both models are deterministic, numerical results are the outcome of a single run.

As stated in the main text, analytical solutions assume weak (*s*, *t* ≪ 1). However, we found that analytical results under relatively strong selection still provided a qualitatively good fit to their corresponding numerical results under both haploid and diploid control of pollen competitive ability (grey series, Figure E1), suggesting interpretations presented in the main text are broadly applicable across a wide range of selection strengths. In Figure E1, weak selection indicates *s* = 0.01 and *t* = 0.03 strong selection indicates *s* = 0.05 and *t* = 0.1.

**Figure E 1.**
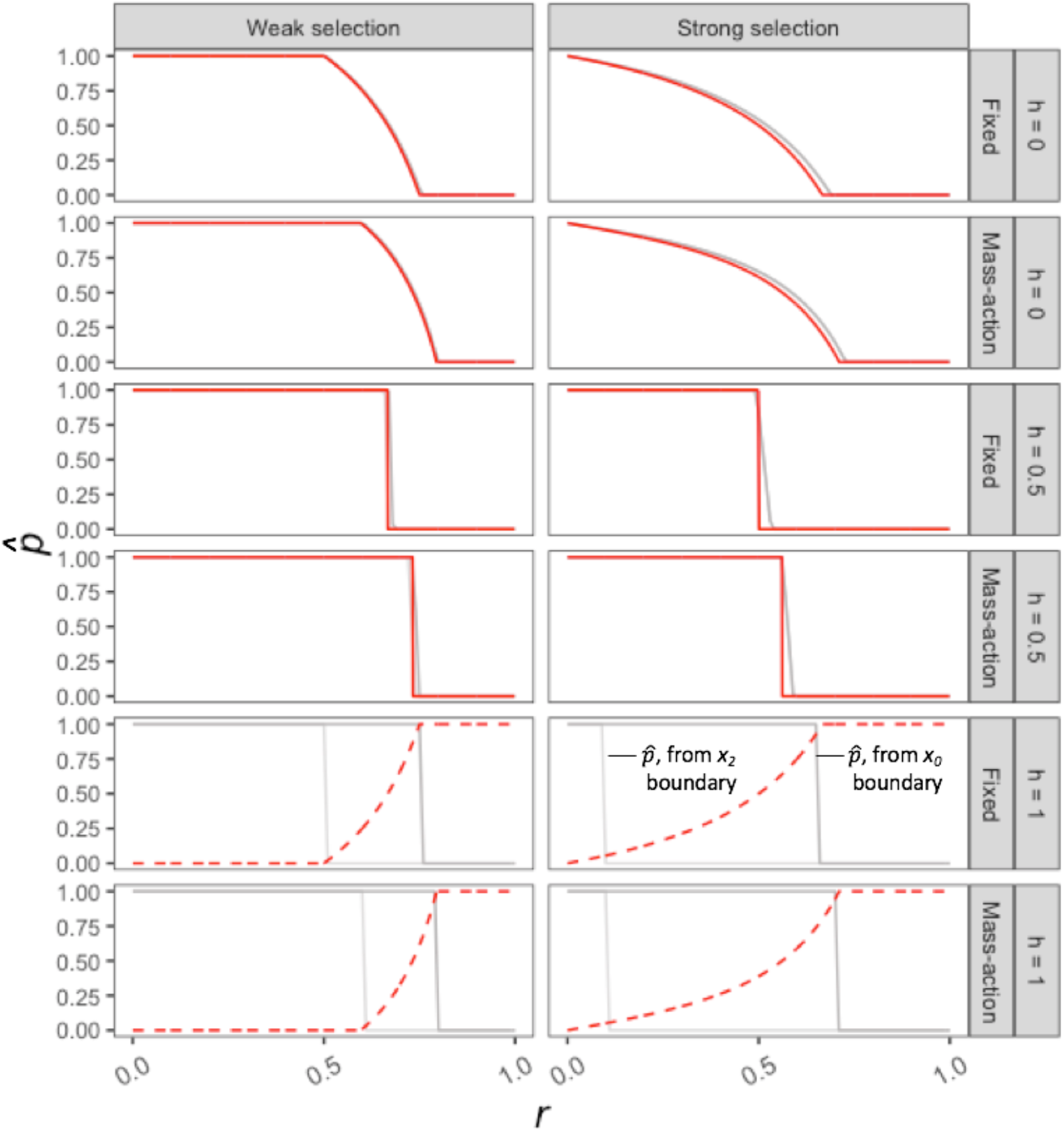

Importantly, when *h* = 1.0, drops in equilibrium allele frequencies from 1 to 0 for all combinations of selection strength and selfing mode are immediate, i.e., there are no intermediate equilibrium values (i.e., 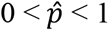). Any slopes are the result of the increment chosen for the *x* axis (0 ≤ *r* ≤ 1 by 0.01).

In addition to a comparison with numerical results under strong selection, we can also compare invasion conditions under fixed selfing assuming weak selection to conditions derived without assuming weak selection. For example, Eqs. 7a and 7b in the main text give maximum and minimum selfing rates for *C_g_* and *C_s_* invasion respectively under weak selection. Without this assumption, these maximum and minimum *r* values are:

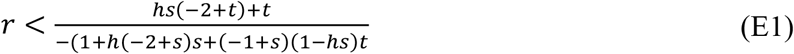

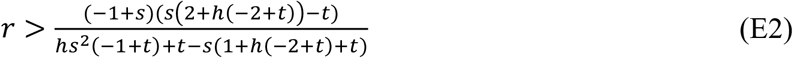

Comparison with expressions 7a and 7b in the main text can be easily considered visually (Figure E2), in which series were generated using *t* = 0.1.

**Figure E 2.**
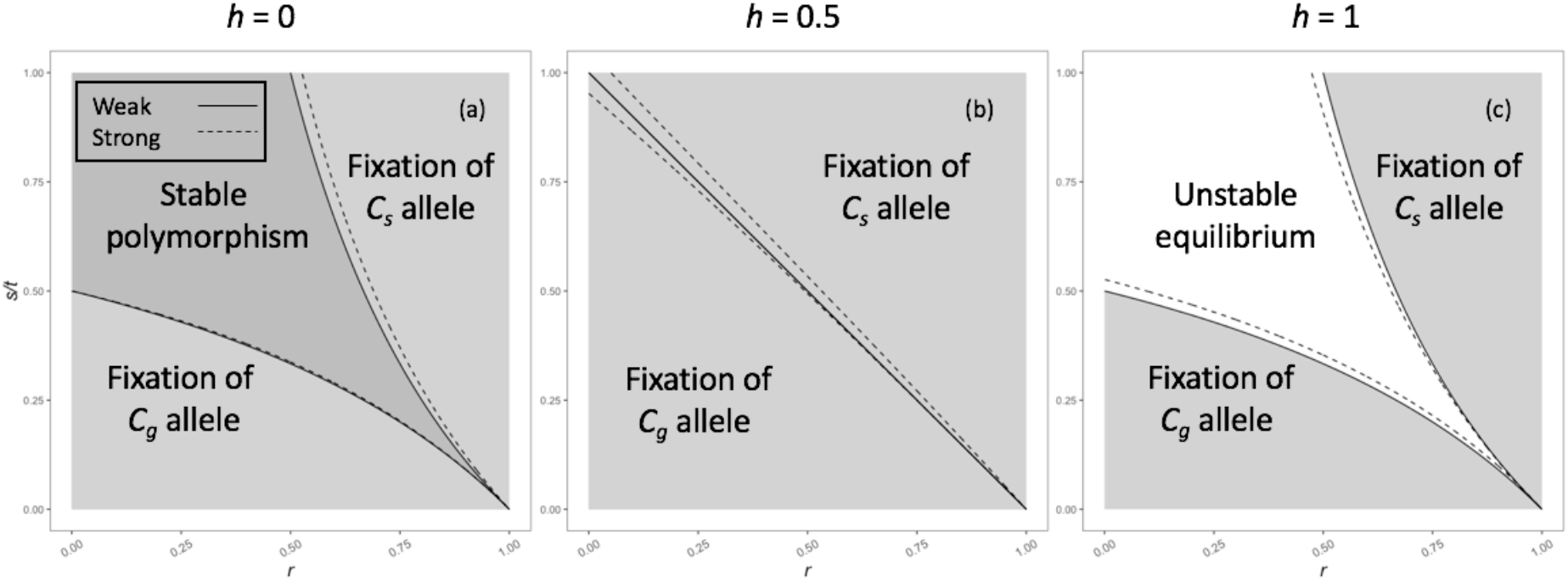

Perhaps most notable in Figure E2 is the small space permitting polymorphism when *h* = 0.5 at low values of *r*. This result is consistent with Figure E1, in which under strong selection there are true numerically-derived intermediate equilibrium allele frequencies (i.e., 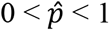) when *h* = 0.5, unlike under weak selection. Overall, we find strong qualitative agreement between invasion conditions predicted both with and without weak selection, suggesting our results likely hold for a broader range of selection parameters than those presented in the main text.

## Appendix F

### Critical selfing rates under mass-action selfing

Using the methods described in the main text, the analytical solutions specifying the selfing rates required for invasion *C_g_* and *C_s_* alleles under mass-action selfing are as follows:

For *C_g_: r* <

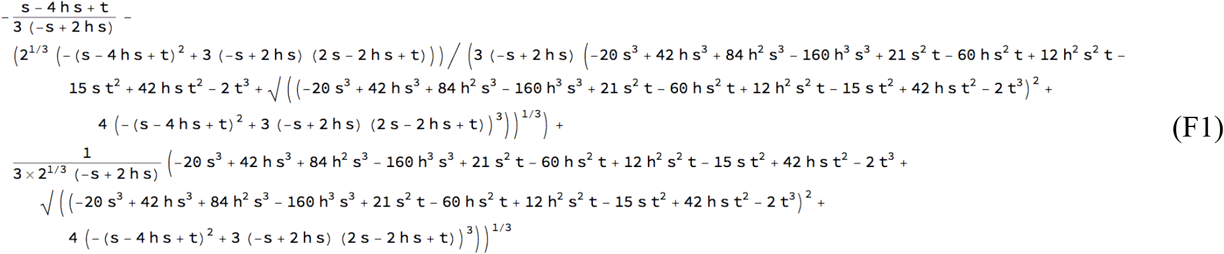

For *C_s_: r* >

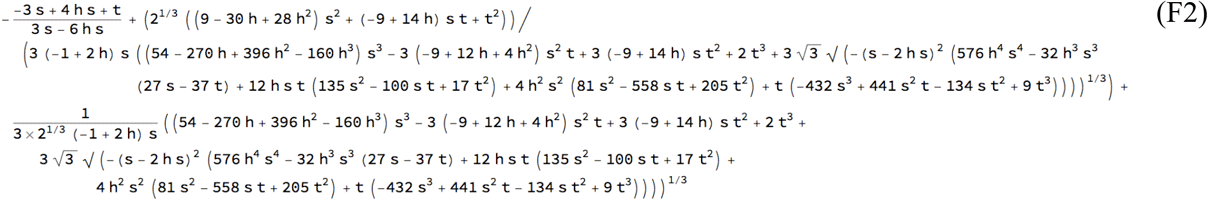

## Appendix G

### Critical diploid selection coefficients under mass-action and fixed selfing

Taking the partial derivative of in-text Eqs. 8a and 8b with respect to *r* produces Eqs. G1 and G2 respectively, which in turn provide insights into the effect of increasing selfing rate on the conditions permitting invasion:

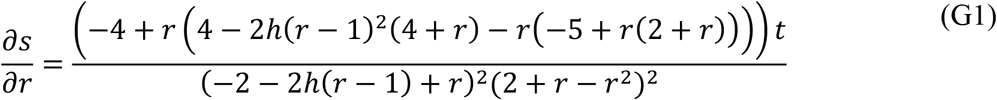

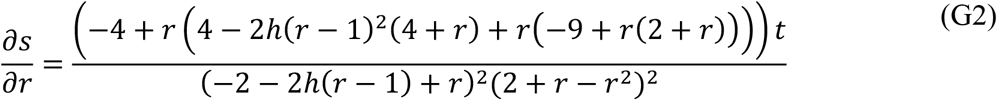

Assuming *h, s, t* and *r* all fall between 0 and 1, the above expressions are always negative, implying the critical value of *s* always decreases as *r* increases (Figure 1). When *h* < 0.5, the maximum value of *s* permitting *C_g_* invasion decreases more slowly as *r* increases than the minimum *s* value permitting *C_s_* invasion. This results in a range of *r* values in which *s* is below the maximum value for *C_g_* invasion and above the minimum value for *C_s_* value, producing the potential for polymorphism. Conversely, when *h* > 0.5, the minimum *s* for *C_s_* invasion decreases more quickly in response to *r* than the maximum value for *C_g_* invasion. This produces a range of *r* values in which neither allele can invade (i.e., unstable equilibrium). Focusing on the effect of selfing on the *C_g_* allele, increasing *r* contracts the parameter space allowing invasion of an allele conferring a benefit to pollen competitiveness.

To compare effects of fixed versus mass-action selfing on invasion, we can compare how minimum and maximum *s* values permitting invasion of *C_s_* and *C_g_* alleles respond to increases in *r* in these two selfing systems. Below are expressions for how the maximum and minimum *s* values change in response to *r* under fixed selfing:

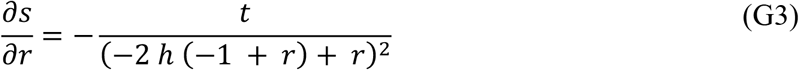

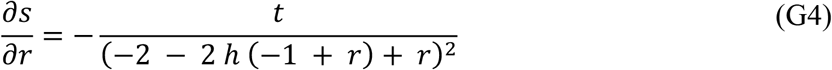

We can easily visualise differences between Eqs. G1,2 and G3,4, and thus how critical *s* values respond to *r* under mass-action versus fixed selfing, by considering the case in which *h* = 0.25 and *t* = 0.03 (Figure G1).

**Figure G 1.**
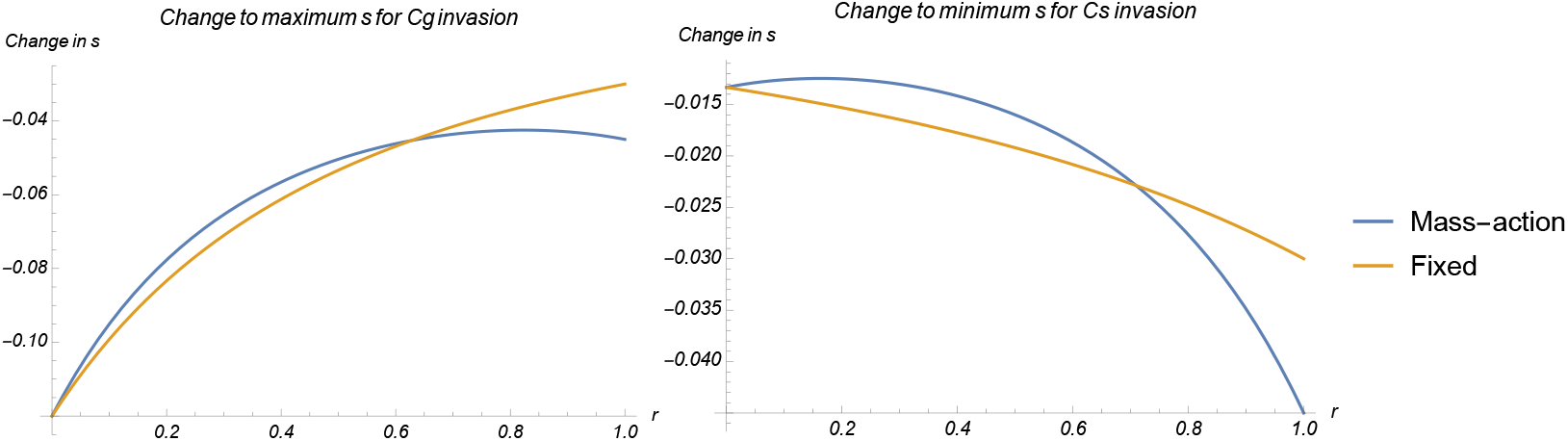

For both maximum and minimum *s* values, it is clear that the critical *s* value decreases more rapidly for fixed selfing than mass-action as *r* increases from 0, as evidenced by the initially lower (*i.e*., more negative) values for the fixed 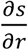 series than for mass-action series. This difference implies that for a given *s* value, the critical *r* value allowing *C_g_* invasion will be lower than that of mass-action selfing. Indeed, we find this to be true.

**Figure G 2.**
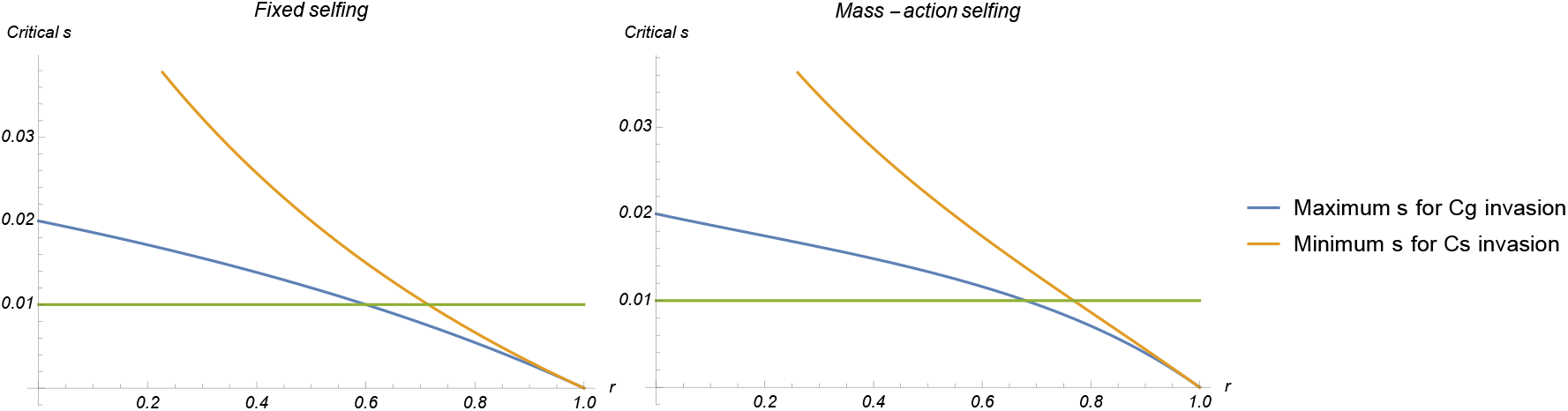

For a given *s* value (in this case 0.01, as given by the green series in Figure G2), both maximum and minimum *s* values permitting invasion are right-shifted along the *r* axis. This indicates the *C_g_* allele can be maintained at higher selfing rates under mass-action selfing than fixed selfing.

Returning to Figure G1, the difference in 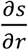 between fixed and mass-action selfing for a given *r* value is greater when considering the maximum *s* permitting *C_g_* invasion than the minimum *s* permitting *C_s_* invasion. While the minimum *s* for *C_s_* invasion is right-shifted under mass-action selfing compared to fixed selfing, the magnitude of this shift is less than that of the corresponding maximum *s* values. Put together, this results in a contracted range of *r* values permitting polymorphism, which can be seen in Figure G2 as the space in which the minimum *s* series falls above the *s* = 0.01 series and the maximum *s* series falls below (0.69 < *r* < 0.77 under mass-action selfing; 0.56 < *r* < 0.73 under fixed selfing).

A similar analysis when *h* > 0.5 yields qualitatively the same results but with unstable equilibria, not protected polymorphism.

## References

Arunkumar, R., E. B. Josephs, R. J. Williamson, and S. I. Wright. 2013. Pollen-specific, but not sperm-specific, genes show stonger purifying selection and higher rates of positive selection than sporophytic genes in *Capsella grandiflora*. Mol. Biol. Evol. 30:2475–2486.

Barrett, S. C. and C. G. Eckert. 2012. 14 Variation and Evolution of Mating Systems in Seed Plants. Biological approaches and evolutionary trends in plants: 229.

Barrett, S. C. H. 1998. The evolution of mating strategies in flowering plants. Trends Plant Sci. 3:335–341.

Baskin, J. M. and C. C. Baskin. 2015. Pollen (microgametophyte) competition: an assessment of its significance in the evolution of flowering plant diversity, with particular reference to seed germination. Seed Sci. Res. 25:1–11.

Borg, M., L. Brownfield, and D. Twell. 2009. Male gametophyte development: a molecular perspective. J. Exp. Bot. 60:1465–1478.

Campbell, S. A. 2015. Ecological mechanisms for the coevolution of mating systems and defence. New Phytol. 205:1047–1053.

Charlesworth, D. and B. Charlesworth. 1992. The effects of selection in the gametophyte stage on mutational load. Evolution 46:703–720.

Crow, J. F. and M. Kimura. 1970. Introduction to Population Genetics Theory. Harper & Row, New York.

Ewing, E. 1977. Selection at the haploid and diploid phases: clinical variation. Genetics. 87:195–208.

Damgaard, C. and V. Loeschcke. 1994. Genetic-variation for selfing rate the dependence of selfing rate on mating history in *Brassica napus* (rape seed). Heredity 72:570–573.

Galloway, L. F. 2001. The effect of maternal and paternal environments on seed characters in the herbaceous plant *Campanula americana* (Campanulaceae). Am. J. Bot. 88:832–840.

Goodwillie, C., S. Kalisz, and C. G. Eckert. 2005. The evolutionary enigma of mixed mating systems in plants: occurrence, theoretical explanations, and empirical evidence. Annu. Rev. Ecol. Evol. Syst. 36:47–79.

Hafidh, S., K. Breznenová, P. Růžička, J. Feciková, V. Čapková, and D. Honys. 2012. Comprehensive analysis of tobacco pollen transcriptome unveils common pathways in polar cell expansion and underlying heterochronic shift during spermatogenesis. BMC Plant Biol. 12(1): 24.

Haldane, J. B. S. 1932. The Causes of Evolution. Reprinted 1990, Princeton University Press.

Harrison, M. C., E.B. Mallon, D. Twell, and R.L. Hammond. 2015. Deleterious mutation accumulation in *Arabidopsis thaliana* pollen genes: a role for a recent relaxation of selection. bioRxiv, 016626.

Holsinger, K. E. 1991. Mass-action models of plant mating systems: the evolutionary stability of mixed mating systems. Am. Nat. 138:606–622.

Igic, B. and J. R. Kohn. 2006. Bias in the studies of outcrossing rate distributions. Evolution 60 (5): 1098–1103.

Immler, S., G. Arnqvist, and S. P. Otto. 2012. f. Evolution 66:55–65.

Jarne, P. and D. Charlesworth. 1993. The evolution of the selfing rate in functionally hermaphrodite plants and animals. Annu. Rev. Ecol. Evol. Syst. 24:441–466.

Jordan, C. Y. and T. Connallon. 2014. Sexually antagonistic polymorphism in simultaneous hermaphrodites. Evolution 68:3555–3569.

Lloyd, D. G. 1979. Some reproductive factors affecting the selection of self-fertilization in plants. The American Naturalist 113(1):67–79.

Mazer, S. J., A. A. Hove, B. S. Miller, and M. Barbet-Massin. 2010. The joint evolution of mating system and pollen performance: Predictions regarding male gametophytic evolution in selfers vs. outcrossers. Perspect. Plant Ecol. Evol. Syst. 12:31–41.

Mazer, S. J., B. T. Hendrickson, J. P. Chellew, L. J. Kim, J. W. Liu, J. Shu, and M. V. Sharma. 2018. Divergence in pollen performance between *Clarkia* sister species with contrasting mating systems supports predictions of sexual selection. Evolution 72(3):453–472.

McCallum, B. and S. M. Chang. 2016. Pollen competition in style: Effects of pollen size on siring success in the hermaphroditic common morning glory, *Ipomoea purpurea*. Am. J. Bot. 103(3):460–470.

Mulcahy, D. L. and G. B. Mulcahy. 1987. The effects of pollen competition. Am. Sci. 75:44–50.

Parker, G. and M. Begon. 1993. Sperm competition games: sperm size and number under gametic control. Proc. R. Soc. London, Ser. B. 253:255–262.

Pélabon, C., L. Hennet, G. H. Bolstad, E. Albertsen, Ø. H. Opedal, R. K. Ekrem, and W. S. Armbruster. 2016. Does stronger pollen competition improve offspring fitness when pollen load does not vary? Am. J. Bot. 103:522–531.

Sarkissian, T. S. and L.D. Harder. 2001. Direct and indirect responses to selection on pollen size in *Brassica rapa* L. J. Evol. Biol. 14(3):456–468.

Schemske, D. W. 1978. Evolution of reproductive characteristics in Impatiens (Balsaminaceae): the significance of cleistogamy and chasmogamy. Ecology 59(3):596–613.

Schemske, D. W. and R. Lande. 1985. The evolution of self-fertilization and inbreeding depression in plants. II. Empirical observations. Evolution 39:41–52.

Smith-Huerta, N. L. 1996. Pollen germination and tube growth in selfing and outcrossing populations of *Clarkia tembloriensis* (Onagraceae). Int. J. Plant Sci. 157:228–233.

Sousa, E., B. Kost, and R. Malhó. (2008). *Arabidopsis* phosphatidylinositol-4-monophosphate 5-kinase 4 regulates pollen tube growth and polarity by modulating membrane recycling. Plant Cell, 20(11), 3050–3064.

Walsh, N. and D. Charlesworth. 1992. Evolutionary interpretations of differences in pollen tube growth rates. Q. Rev. Biol. 67:19–37.

Winn, A. A., & Moriuchi, K. S. 2009. The maintenance of mixed mating by cleistogamy in the perennial violet *Viola septemloba* (Violaceae). American Journal of Botany 96(11):2074–2079.

Winsor, J., L. Davis, and A. Stephenson. 1987. The relationship between pollen load and fruit maturation and the effect of pollen load on offspring vigor in *Cucurbita pepo*. Am. Nat. 129:643–656.

Winsor, J. A., S. Peretz, and A. G. Stephenson. 2000. Pollen competition in a natural population of *Cucurbita foetidissima* (Cucurbitaceae). Am. J. Bot. 87:527–532.

Young, H. J. and M. L. Stanton. 1990. Influence of environmental quality on pollen competitive ability in wild radish. Science 248:1631–1633.

